# Simultaneous adjustments of major mitochondrial pathways through redox regulation of dihydrolipoamide dehydrogenase (mtLPD1)

**DOI:** 10.1101/2022.04.02.486831

**Authors:** Stefan Timm, Nicole Klaas, Janice Niemann, Kathrin Jahnke, Saleh Alseekh, Youjun Zhang, Paulo V.L. Souza, Liang-Yu Hou, Peter Geigenberger, Danilo M. Daloso, Alisdair R. Fernie, Martin Hagemann

## Abstract

Thioredoxins (TRX) are pivotal for the redox regulation of enzyme activities to adjust metabolic fluxes towards environmental changes. Previous reports demonstrated TRX *o1* and *h2* impact on mitochondrial metabolism including photorespiration and the tricarboxylic acid (TCA) cycle. Here, we aimed to unravel potential specificities between regulation modes of both TRXs, especially under conditions with short-term changes in photorespiration. Therefore, short-term metabolite responses of single *TRX* mutants were analyzed after exposure to altered CO_2_/O_2_ ratios during darkness and illumination. This approach was complemented by comprehensive characterization of multiple Arabidopsis mutants lacking either one or both *TRX* in the wild-type Arabidopsis or the glycine decarboxylase (GDC) T-protein knock down line (*gldt1*). The results provided evidence for additive effects of combined TRX *o1* and *h2* deficiency to suppress growth, photosynthesis and mitochondrial metabolism. Quantification of pyrimidine nucleotides in conjunction with metabolite and ^13^C-labelling approaches revealed a rather uniform impact on mitochondrial dihydrolipoamide dehydrogenase (mtLPD1) dependent pathways. Biochemical analysis of recombinant mtLPD1 demonstrated its inhibition by NADH, pointing at an additional measure to fine-tune it’s *in vivo* activity. Collectively, we propose that TRX *o1* and *h2* contribute to the communication of altered subcellular redox-states through direct and indirect regulation of mtLPD1. This regulation module might represent a common intercept for simultaneous adjustments in the operation of photorespiration, the TCA-cycle and the degradation of branched chain amino acids.

**One-sentence summary:** Redox regulation of mitochondrial dihydrolipoamide dehydrogenase (mtLPD1) simultaneously modulates photorespiration, the tricarboxylic acid (TCA)-cycle and branched chain amino acid (BCAA) degradation in response to rapid environmental changes.

## Introduction

Thioredoxins (TRX) catalyze reduction-oxidation (redox) reactions via reversible thiol-disulfide exchanges in numerous enzymes of primary metabolism in many organisms. This posttranslational modification (PTM) is key to maintain fast and flexible adjustments of metabolism towards fluctuating conditions within seconds to hours. The possibility to allow efficient acclimation of metabolism is of particular significance for plants as they are sessile and unable to actively escape from rapid and harsh environmental changes that frequently occur in nature. The latter have been proposed to lead to strong alterations in plant subcellular redox states, which requires redox regulation at multiple levels (Zaffagnini et al., 2019). Regulation via TRX is mainly adjusting enzymatic activities and, in turn, metabolic fluxes. However, this regulatory principle is of broad importance also for many fundamental processes in plants including light acclimation, stress tolerance, gene expression, protein transport and processing, autophagy, seed germination and plant development (e.g. Balmer et al., 2004; Buchanan, 2016; Geigenberger et al., 2017; Lee et al., 2021; Møller et al., 2020; Nietzel et al., 2020).

Within the last years it was demonstrated that plants possess a complex TRX network. For example, *Arabidopsis thaliana* L. (Arabidopsis) contains seven different TRX families, encoded by at least 20 different genes, distributed among various subcellular compartments. The best studied TRX are isoforms belonging to *m*, *f*, *x*, *y* and *z* types that exclusively localize to the chloroplast. Within this compartment, they are involved in various processes including the regulation of light acclimation, photosynthesis, starch metabolism, as well as carbon and nitrogen utilization (e.g., Ancín et al., 2021; Buchanan, 2016; Geigenberger et al., 2017; Nikkanen et al., 2017; Thormählen et al., 2013; 2017). Important regulatory roles have also been shown for extra-plastidal TRX belonging to the *o* and *h* types. Whilst TRX *o* isoforms are restricted to mitochondria and, perhaps, the nucleus, the TRX *h* isoforms can be found in diverse subcellular compartments (Delorme-Hinoux et al., 2016, Buchanan, 2017; Lee et al., 2021). Daloso et al (2015) demonstrated that TRX *o1* is involved in the regulation of the TCA-cycle through controlling the activity of succinate dehydrogenase (SDH) and fumarase (FUM), thereby contributing to the light-inactivation of respiration and the establishment of the different flux-modes in the TCA-cycle (Sweetlove et al., 2010; Daloso et al., 2015; daFonseica-Pereira et al., 2020). Moreover, TRX *o1* was discussed to contribute to regulation of mitochondrial alternative oxidase (AOX) under certain conditions but its significance *in vivo* remains controversial (Florez-Sarasa et al., 2019; Møller et al., 2020). Cumulative evidence, however, exists for TRX *o1* mediated regulation of photorespiration. It was demonstrated that TRX *o1* deficiency affects mitochondrial photorespiratory metabolism in response to shifts from high-to-low CO_2_ and, perhaps, light-induction of photosynthesis (Reinholdt et al., 2019a; 2019b). In addition, biochemical analysis revealed the direct involvement of TRX *o1* in the redox regulation of mitochondrial dihydrolipoamide dehydrogenase (mtLPD1). It is worth mentioning that similar observations were reported with TRX *h2*, a member of the largest TRX family in Arabidopsis (daFonseica-Pereira et al., 2020). Although a few reports suggested mitochondrial localization of TRX *h2*, a recent study revealed its localization to the microsomal fraction (Hou et al., 2021). Nevertheless, there is experimental evidence that TRX *h2* deficiency impacts mitochondrial photorespiratory metabolism, while the underlying mechanism remained elusive to date (daFonseica-Pereira et al., 2020; 2021). It seems tempting to hypothesize that TRX *h2* operation also contributes to the mitochondrial redox-homeostasis. A likely central target could be glycine decarboxylase (GDC), because TRX *h2* mutant lines showed altered photorespiration under sudden environmental changes. This assumption is supported by biochemical analysis showing changed GDC activities under redox-changes. In detail, GDC activity as a whole was shown to be inhibited at high NADH/NAD^+^ ratios, whilst its activity can be modulated also through redox regulation of the single GDC proteins P and mtLPD1 (Bourguignon et al., 1988; Hasse et al., 2013; Reinholdt et al., 2019a, daFonseica-Pereira et al., 2020).

The findings that TRX *o1* and *h2* are able to redox regulate mtLPD1 eventually has consequences beyond photorespiration, since mtLPD1 is not restricted to GDC. Indeed, the protein is also shared with two TCA-cycle complexes, namely, 2-oxoglutarate dehydrogenase (OGDC) and pyruvate dehydrogenase (PDC), as well as the branched chain 2-oxoacid dehydrogenase complex (BCKDC), involved in the catabolism of branched chain amino acids (BCAAs) (Bourguignon et al., 1988; Millar et al., 1998; 1999). Consequently, mtLPD1 might be a prime candidate for the simultaneous regulation of three major branches of mitochondrial metabolism towards environmental changes as suggested recently (Timm and Hagemann, 2020). To test this hypothesis, we compared the single *TRX o1* (*trxo1-1*) and *h2* (*trxh2-2*) mutants with double and triple Arabidopsis mutants lacking *TRX o1* and *h2* in the wild-type or in the *gldt1-1* mutant background (Engel et al., 2008). The mutant *gldt1* was included, because it has a moderate reduction in the photorespiratory flux through a knock down in the GDC T-protein expression, decreasing overall GDC activity (Timm et al., 2018). The subsequent analyses focused on photorespiration, the TCA-cycle and the degradation of BCAAs. Additionally, we also compared short-term GDC operation in response to onset of illumination with active and suppressed photorespiration using metabolomics and an ^13^C-isotope labelling experiment. Finally, the mutant analysis was complemented by continuative biochemical analysis of recombinant mtLPD1. The results obtained point to mtLPD1 as a potential master switch translating alterations in the mitochondrial redox-state to modulate flux through photorespiration, the TCA-cycle and the degradation of BCAAs in response to rapid environmental changes involving TRX *o1* and *h2*.

## Results

### Targeted analysis of mtLPD1-dependent pathways in single *TRX* mutants exposed to short-term environmental changes

Previous reports showed deletion of *TRX o1* and *h2* affect the performance of mitochondrial pathways including photorespiration (Reinholdt et al., 2019a; daFonseica-Pereira et al., 2020) the TCA-cycle (Daloso et al., 2015) and, perhaps, BCAAs degradation (Timm and Hagemann, 2020). Hence, single *TRX o1* or *h2* deficiency mutants were initially analyzed whether they show similar or specific metabolic responses among the three mtLPD1-dependent mitochondrial pathways, particularly under conditions with short-term changes in photorespiration. To this end, the wild type was grown next to the single *TRX* mutants (*trxo1-1* and *trxh2-2*) under standard conditions and then exposed for 15 min to altered CO_2_/O_2_ ratios in the absence and presence of light (Fig. 1A). Subsequently, selected metabolite amounts were quantified through liquid chromatography coupled to tandem mass spectrometry (LC-MS/MS). Glycine, the photorespiratory intermediate metabolized by the GDC in mitochondria, showed low concentrations under conditions with no or suppressed photorespiration. Furthermore, glycine was largely invariant among the wild type and the *TRX* single mutants. Under photorespiratory conditions, however, glycine is generally present in higher amounts in all genotypes. Significantly higher glycine accumulation was observed in both *TRX* mutants compared to the wild type at enhanced photorespiration, i.e. at 21 and 40% O_2_ and 500 µmol m^-2^ s^-1^ light (Fig. 1B). Serine concentrations are almost similar in all conditions and do not show much variations between the wild type and the *TRX* mutants except for a slight but significant increase in darkness at 3% O_2_ (Fig. 1B). The BCAAs pattern is similar to that of glycine. Almost no significant changes appeared in the absence or with suppressed photorespiration (except decreases in leucine in *trxo1* and *trxh2* and isoleucine in *trxo1*) among the genotypes (Fig. 1C). After induction of photorespiration all three BCAAs accumulated in the *TRX* mutants at 150 µmol m^-2^ s^-1^ light in the presence of 21 and 40% O_2_. These changes were more distinct at the higher light intensity of 500 µmol m^-2^ s^-1^ in conjunction with 21 and 40% O_2_ (Fig. 1C).

**Figure 1.**
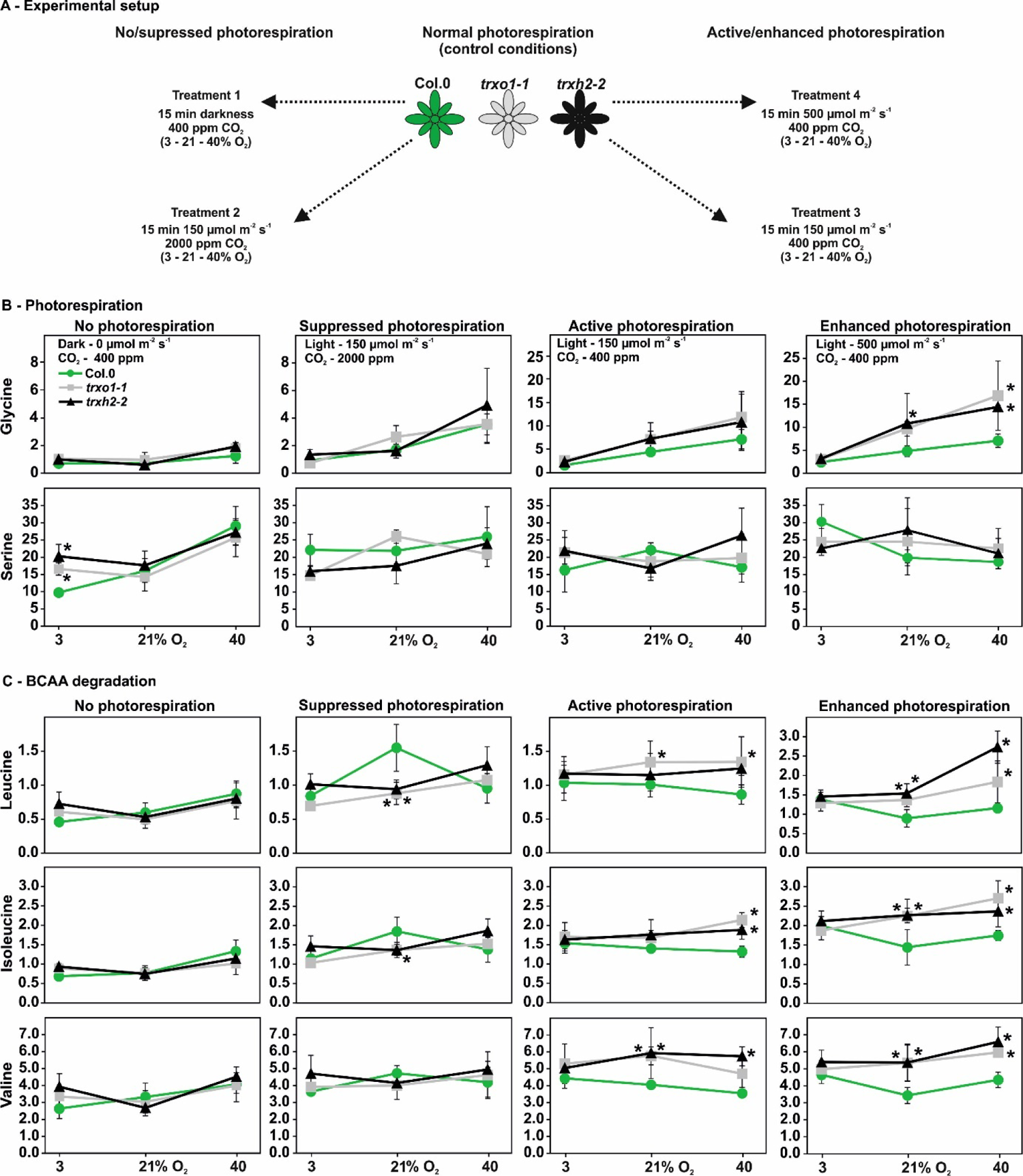
Experimental setup and amounts of selected intermediates related to photorespiration and BCAA degradation in single *TRX* mutants and the wild type. **(A)** Overview of the experimental setup. Plants were grown under standard conditions (150 µmol m^-2^ s^-1^ light, 400 ppm CO_2_, 21% O_2_) to stage 5.1 (Boyes et al., 2001) and subsequently used for short-term environmental treatments as follows: (I) no photorespiration – 0 µmol m^-2^ s^-1^ light (darkness), 400 ppm CO_2_, 3, 21 and 40% O_2_; (II) suppressed photorespiration – 150 µmol m^-2^ s^-1^ light, 2000 ppm CO_2_, 3, 21 and 40% O_2_; (III) active photorespiration - 150 µmol m^-2^ s^-1^ light, 400 ppm CO_2_, 3, 21 and 40% O_2_; and (IV) enhanced photorespiration - 500 µmol m^-2^ s^-1^ light, 400 ppm CO_2_, 3, 21 and 40% O_2_, respectively. For all treatments we used plants in the second half of the illumination phase (4 to 10 h light) to ensure fully active and stable photosynthesis. Absolute metabolite amounts (nmol mg^-1^ FW^-1^) were quantified by LC-MS/MS using leaf-discs (2 cm^-2^) harvested after 15 minutes exposure to each condition following attachment of fully expanded leaves to the Licor chamber. Shown are mean values ± SD (n = 4) of **(B)** selected photorespiratory intermediates and **(C)** branched chain amino acids. Asterisks indicate significant alterations of the *TRX o1* and *h2* single mutant compared with the wild type in each condition according to Student’s *t*-test (\**p* < 0.05).

A more diverse picture was observed among the TCA-cycle intermediates. In the absence of photorespiration, a consistent pattern was observed in both *TRX* mutants for most intermediates including pyruvate, citrate, aconitate, 2-oxoglutarate and succinate. Almost all of these intermediates were elevated at 3% O_2_ (except succinate) and decreased at 21 and 40% O_2_ (except pyruvate and 2-oxoglutarate). Fumarate and malate, however, were largely invariant and only elevated in *trxo1* at 40% O_2_ (Fig. 2). In response to illumination, with suppressed photorespiration, fewer changes emerged. Almost no systematic changes were found in pyruvate, 2-oxoglutarate, fumarate and malate. Increases were seen in succinate at 3% O_2_ (both lines), citrate at 3% O_2_ (*trxh2*) and 21% O_2_ (both lines) and aconitate at all O_2_ concentrations and both *TRX* mutants (except *trxo1* at 3%). Under conditions with active photorespiration and standard growth light, pyruvate and citrate remained unaltered among all genotypes. Most of the other intermediates did show only few, non-systematic changes. Interestingly, aconitate and fumarate displayed a somewhat reversed accumulation pattern between *trxo1* and *trxh2*. Finally, in response to a higher light intensity both *TRX* mutants significantly accumulated TCA-cycle intermediates at one or both higher O_2_ concentrations including pyruvate, citrate, aconitate, succinate fumarate and malate. A slight decrease of 2-oxoglutarate was seen in *trxh2* at 40% O_2_ (Fig. 2). Collectively, single knock outs of the TRX *o1* and *h2* have similar effects on photorespiratory glycine and BCAAs under conditions of elevated photorespiration, whereas the effects on TCA cycle intermediates were less homogenous.

**Figure 2.**
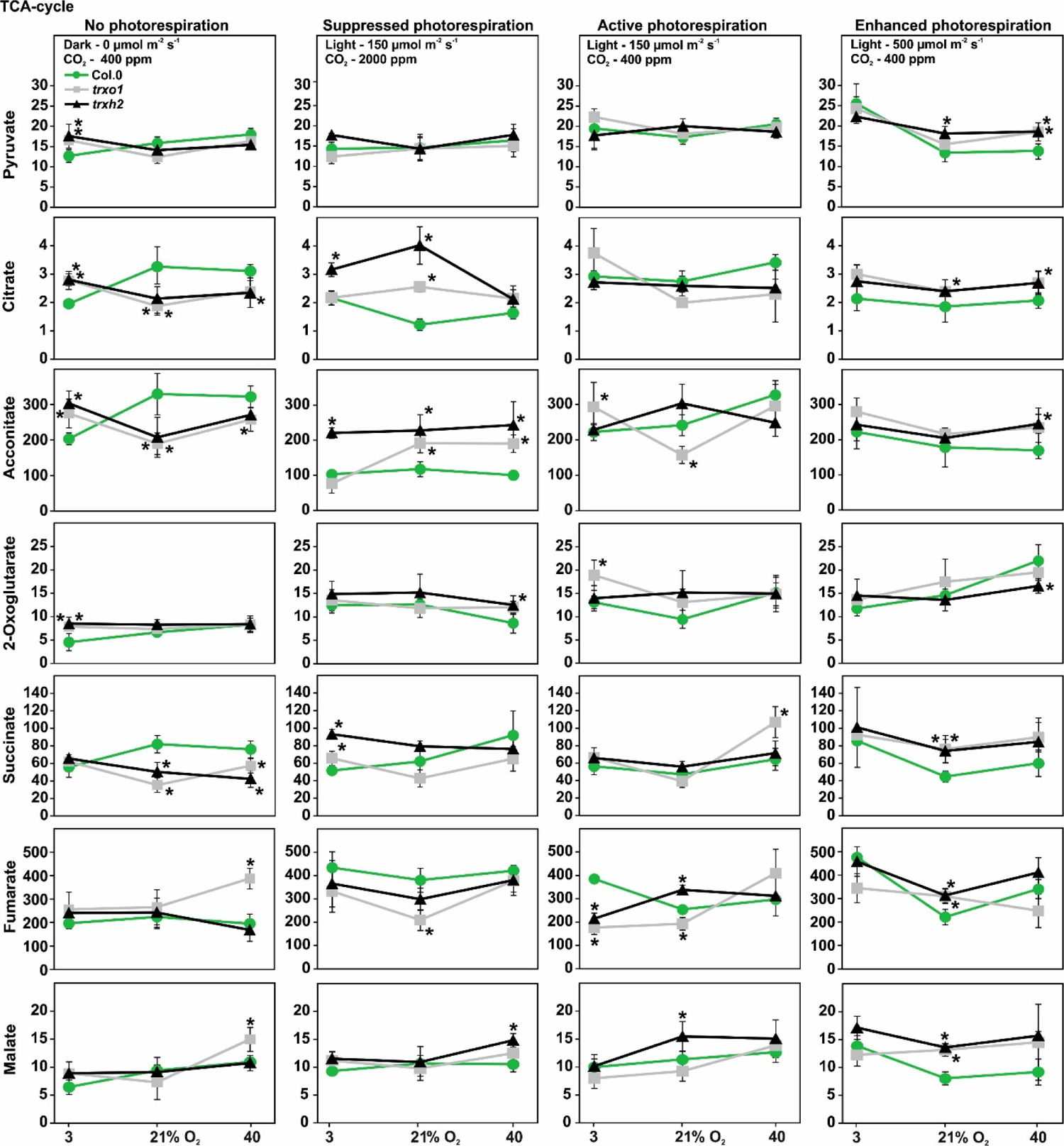
Amounts of TCA-cycle intermediates in single *TRX* mutants and the wild type. Sampling and metabolite analysis were carried out as described in the legend of Fig. 1. Shown are mean values ± SD (n = 4) of selected TCA-cycle intermediates. Asterisks indicate significant alterations of the *TRX o1* and *h2* single mutant compared with the wild type in each condition according to Student’s *t*-test (\**p* < 0.05).

### Combined deletion of *TRX o1* and *h2* additively impairs photosynthetic-photorespiratory gas exchange

Single mutants defective in either, *TRX o1* or *h2*, respectively, do not display major growth retardations or impairments of photosynthetic CO_2_ assimilation if grown under standard conditions with shorter photoperiods. However, an increase in the stoichiometry of photorespiratory CO_2_ release was seen in both single *TRX* mutants during oxygen-dependent gas exchange measurements (Reinholdt et al., 2019a; daFonseica-Pereira et al., 2020). To answer the question, if TRX *o1* and *h2* are redundant or fulfil specific roles, a mutant bearing a combined deletion of *TRX o1* and *h2* was generated (*trxo1h2*; Hou et al., 2021) and specifically analyzed regarding photorespiration as well as mitochondrial metabolism during this study. For this purpose, *trxo1h2* was grown next to the wild type in ambient air (400 ppm CO_2_) to growth stage 5.1 (Boyes et al., 2001) and used for gas exchange measurements. Whilst single *trxo1* and *trxh2* mutants displayed wild-type-like net CO_2_ uptake rates (*A*) in normal air (Reinholdt et al., 2019a; daFonseica-Pereira et al., 2020), the double mutant already showed a significant decrease of *A* to about 15% under these conditions (Fig. 3A). At elevated photorespiratory pressure (50% O_2_), this decrease was further pronounced (up to ∼19%), resulting in an oxygen inhibition of A to about ∼32% (Fig. 3B). In contrast, if the photorespiratory pressure was reduced (3% O_2_ or 2000 ppm CO_2_), *A* values were statistically invariant between the double mutant and the control (Fig. 3A, 3C). Consistent observations were made regarding the CO_2_ compensation point (Γ), i.e. invariant among the genotypes at 3% O_2_, but gradually decreased in the double mutant at 21% and 50% O_2_ compared to wild type (Fig. 3D). These changes lead to an increase of the slope (γ) from the estimated Γ-versus-oxygen response lines by about ∼28% (Fig. 3E), indicating a higher fraction of CO_2_ released from photorespiration. Furthermore, the determination of *A* at increasing light intensities revealed a stronger reduction of photosynthetic CO_2_ uptake rates in *trxo1h2* compared to the wild type most notable at high light (Fig. 3F).

**Figure 3.**
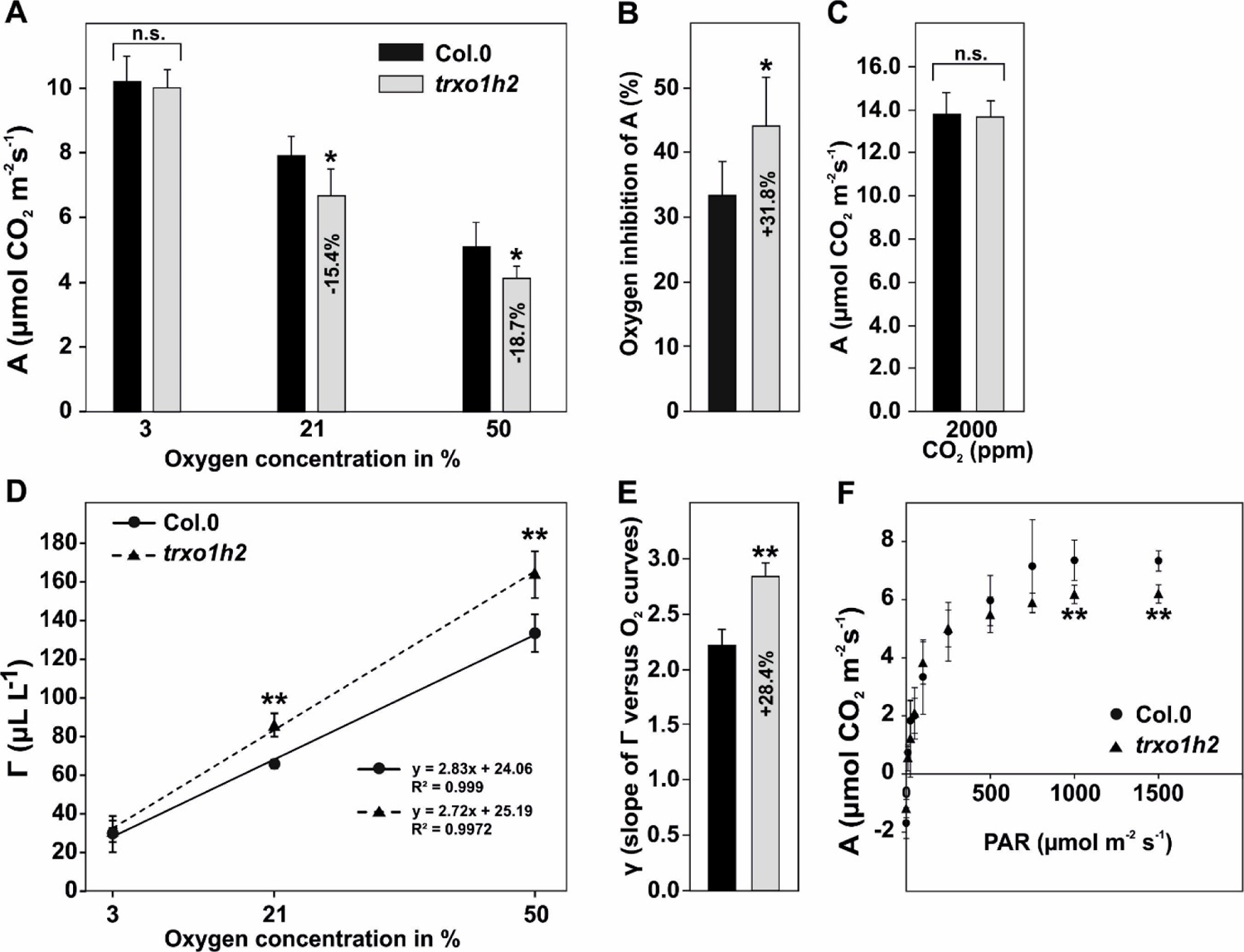
O_2_-dependet gas exchange of the wild type and the *trxo1h2* mutant. Photosynthetic gas exchange parameters at varying O_2_ concentrations (3%, 21%, and 50% O_2_, balanced with N_2_), were determined from plants grown for 8 weeks in normal air (10/14 h day-/night-cycle, 390 ppm CO_2_) to growth stage 5.1 (Boyes et al., 2001). Shown are mean values ± SD (n > 5) of **(A)** net CO_2_ uptake rates (A) at 390 ppm CO_2_, **(B)** oxygen inhibition of A, **(C)** net CO_2_ uptake rates (A) at 2000 ppm CO_2_, **(D)** CO_2_ compensation points (Γ) and **(E)** slopes of the Γ-versus-O_2_ concentration curves (γ). **(F)** Net CO_2_ uptake rates (*A*) at different light intensities. Asterisks indicate significant alterations of the *trxo1h2* mutant compared with the wild type according to Student’s *t*-test (\**p* < 0.05, \*\**p* < 0.01, n. s. - not significant).

The above findings indicate that the TRX *o1* and *h2* are not necessarily redundant but rather have additive impact on photosynthetic gas exchange. The suggested additive impairment of photorespiration in *trxo1h2*, i.e. increased O_2_ sensitivity, is provided by the phenotypic alterations of the *trxho1h2* double mutant under conditions requiring higher photorespiratory capacities. Following growth under a 10/14 h day-/night-cycle in HC and a transfer to LC combined with long day conditions, *trxo1h2* displayed strongly retarded growth compared to all other genotypes and to continuous growth in shorter photoperiods. In contrast, the single *trxo1* or the *trxh2* single deletions are visually indistinguishable from the wild type (Supp. Fig. S1). Furthermore, onset of flowering was comparable between the wild type and the single mutants; however, it was delayed to about 10-to-14 days in *trxo1h2* under this growth condition.

### Distinct accumulation of diagnostic photorespiratory intermediates in *trxo1h2* at high O_2_

Next, the reduced photorespiratory capacity in *trxo1h2* was verified on the metabolite level. To this end, three diagnostic photorespiratory intermediates were quantified via LC-MS/MS in response to short-term high O_2_ and light treatments to stimulate RuBP oxygenation and subsequent photorespiration. 2-Phosphoglycolate (2PG) was selected as a proxy for ribulose-1,5-bisphosphate (RuBP) oxygenation, whilst glycine and serine served as indicators for GDC activity in mitochondria. As expected, the wild type displayed increased amounts of 2PG and glycine after exposure to 40% O_2_ for 15 min at 150 µmol m^-2^ s^-1^ light (Fig. 4). This response was further amplified with a simultaneous increase in the light intensity to 500 µmol m^-2^ s^-1^. Despite serine amounts were largely invariant in all conditions, the glycine-to-serine ratio increased in response to higher O_2_ and light quantities (Fig. 4). The *trxo1h2* mutant showed elevated 2PG (+87%) and serine (+30%) but not glycine contents under control conditions compared to wild type. However, stimulation of RuBP oxygenation and photorespiration though an increased O_2_ concentration (40%) caused much higher accumulation of 2PG (+105%) and glycine (+172%) in the *trxo1h2* double mutant than in wild type. This effect was further amplified with a higher light intensity (Fig. 4). Despite serine appeared wild-type-like at both light intensities and elevated O_2_, the alterations in glycine caused a significant increase in the glycine-to-serine ratio in *trxo1h2* (Fig. 4).

**Figure 4.**
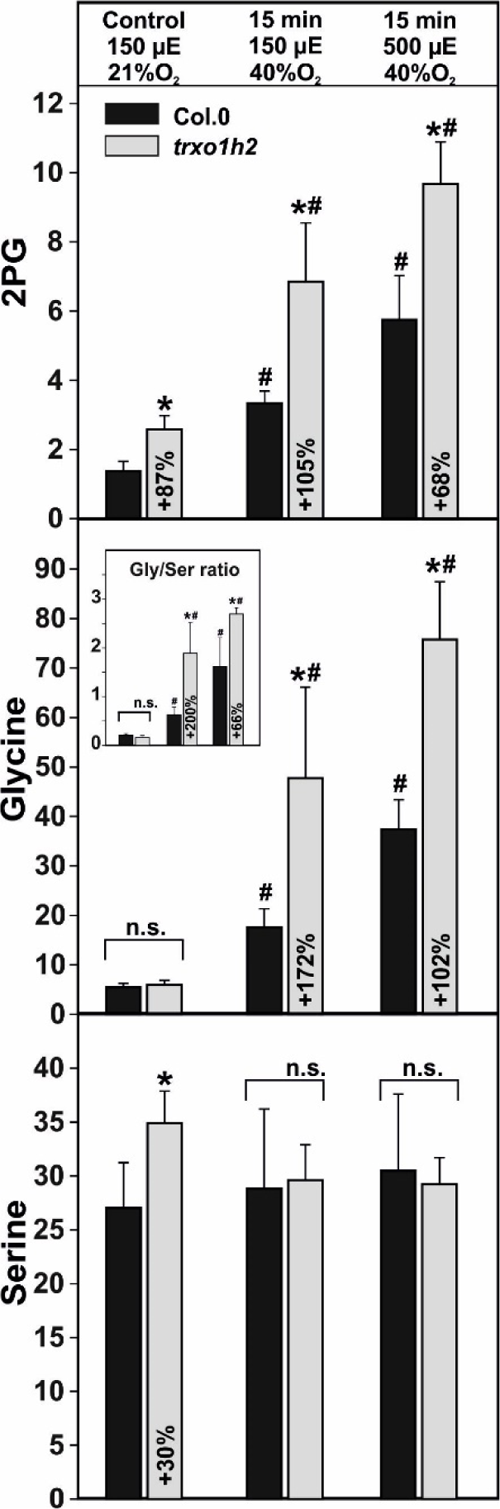
Diagnostic photorespiratory intermediates in the wild type and *trxo1h2*. Absolute metabolite amounts (nmol mg^-1^ FW^-1^) were quantified by LC-MS/MS analysis from plants grown for 8 weeks in normal air (10/14 h day-/night-cycle, 390 ppm CO_2_) to growth stage 5.1 (Boyes et al., 2001). At this stage, fully expanded plant-leaf’s were attached to the Licor chamber in the second half of the illumination phase (4 to 10 h light) for controlled manipulation of O_2_ and light conditions. Leaf-discs (2 cm^-2^) were harvested under growth conditions (control, 21% O_2_, 150 µE light) and after 15 min at 40% O_2_ under 2 different light intensities (150 or 500 µE). Shown are mean values ± SD (n = 4). Asterisks indicate significant alterations of *trxo1h2* compared with the wild type in each condition and rhombs to the control condition according to Student’s *t*-test (\**p* < 0.05, ^#^*p* < 0.05, n.s. - not significant).

### Isolation, phenotype and PSII quantum yield of multiple *TRX/GDC-T* mutants

We previously demonstrated that deletion of *TRX o1* in the *gldt1* mutant knockdown background strengthened the phenotypic and physiological alterations of both single mutants (Reinholdt et al., 2019a). Therefore, the *gldt1* mutant was used to generate *TRX h2* deficiency that permits the comparison of the *trxh2*x*gldt1* and the *trxo1*x*gldt1* double mutants. Furthermore, the triple mutant, simultaneously lacking *TRX o1* and *h2* in the *gldt1* knockdown background, was obtained to clarify if additive effects occur through absence of both TRX. To achieve these goals, the *trxo1*x*gldt1* double mutant (Reinholdt et al., 2019a) was crossed with the single *trxh2-2* mutant allele. After verification of the triple heterozygous T1-intermediate, the T2-generation was grown under high CO_2_ conditions (HC, 3000 ppm CO_2_) to ensure isolation of mutants strongly impaired in photorespiration. Indeed, this procedure led to isolation of a *trxh2*x*gldt1* double and a *trxo1*x*gldt1*x*trxh2* (triple) mutant (Supp. Fig. S2).

The full mutant set was analyzed regarding selected phenotypic and photosynthetic parameters. In detail, the growth and PS efficiency (*F_v_*/*F_m_*) of the genotypes was compared when grown in either condition with suppressed photorespiration (HC - high CO_2_, 3000 ppm) or active photorespiration (LC – low CO_2_, ambient air, 390 ppm). The *adg1-1* mutant, deficient in the small subunit of ADP-glucose pyrophosphorylase (Lin et al., 1988), was included as an additional control, since it displays visible growth retardation under the conditions used which cannot be complemented by elevated CO_2_. All genotypes, except for *adg1*, grew similar to the wild type in HC (Fig. 5A). This visual impression was corroborated by invariant t rosette diameters and total leaf-counts (again with the exception of *adg1*) (Table 1). Similarly, chlorophyll (Chl) a fluorescence images and the *F_v_*/*F_m_* values of the genotypes did not show major differences. Exceptionally, *adg1* displayed a slight reduction in the maximum quantum yield of PSII (Fig. 5A, Table 1). By contrast, distinct changes in growth and Chl a fluorescence emerged during growth in ambient air. Visually, there were no major growth changes between the wild type and the single *TRX* mutants (Fig. 5B). However, the *TRX* double mutant was significantly reduced in its diameter and leaf-number in ambient air, whilst *gldt1* also showed the expected reduction in growth (Fig. 5B, Table 1). More obvious, both double mutants in the *gldt1* background were considerably more retarded in growth compared to *gldt1*, even including yellowish bleaching leaves. Interestingly, the effects were somewhat stronger in the *trxh2*x*gldt1* compared to the *trxo1*x*gldt1*. Nevertheless, all alterations were further amplified in the triple mutant. This statement agreed well with reduced *F_v_*/*F_m_*, rosette diameters and the total leaf-count in air (Fig. 5B, Table 1). Collectively, the phenotypes of the different *TRX* mutations in the *gldt1* background support the notion that deletion of both TRX caused additional effects under ambient conditions in Arabidopsis, at least to some extent.

**Figure 5.**
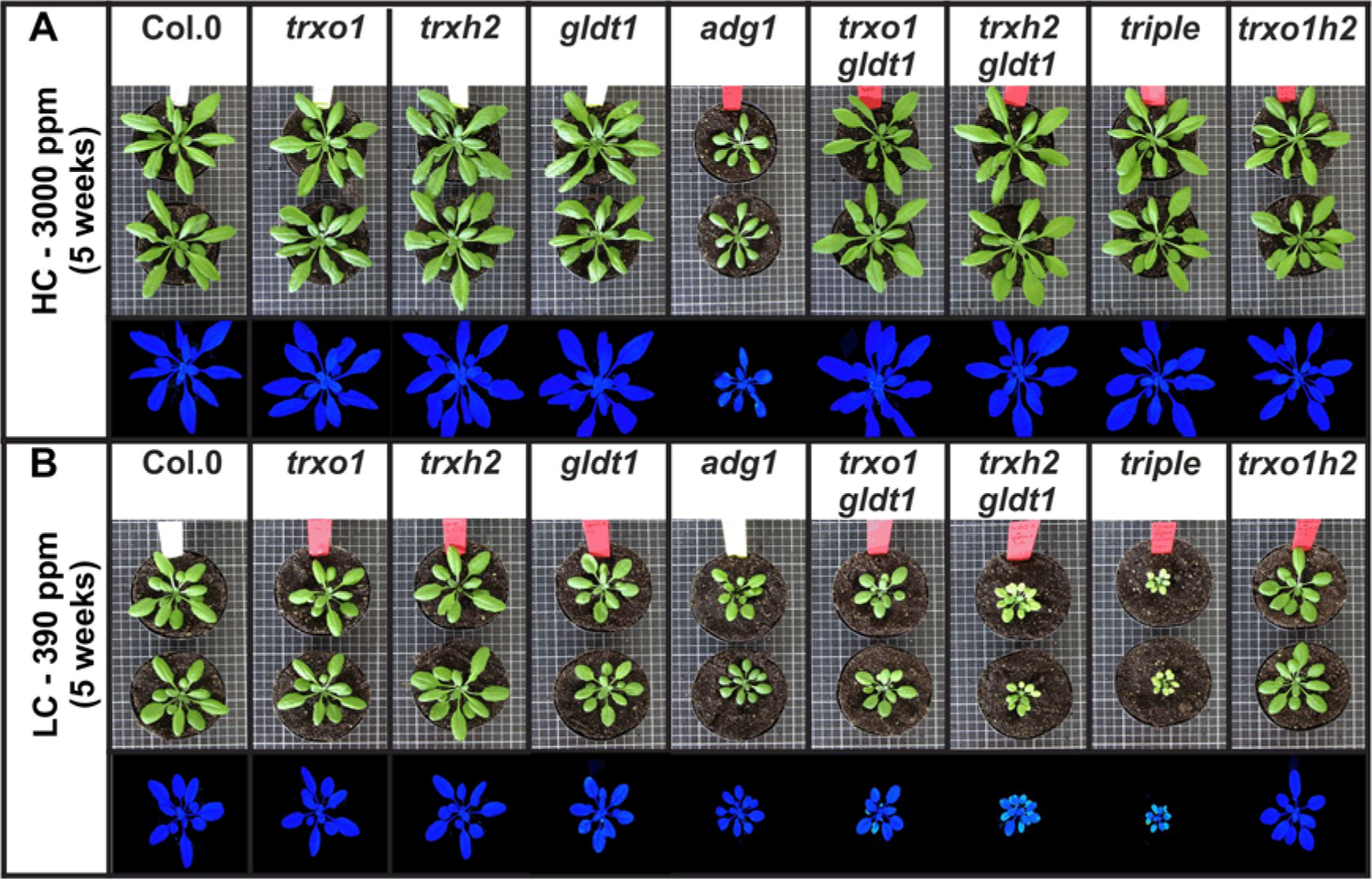
Phenotype and chl a fluorescence images of the mutant set. Plants were grown under environmental controlled conditions (Percivall, 10 h/14 h day-/night-cycle) for 5 weeks in **(A)** high CO_2_ (HC) and **(B)** low CO_2_ (LC, normal air) conditions. At this time point, representative photographs were taken and chl a fluorescence measurement carried out. For quantitative data see Table 1.

**Table 1.** Growth and *F_v_*/*F_m_* of the mutant set and the wild type in HC and LC conditions. Plants were grown under environmental controlled conditions in HC (3000 ppm CO_2_) or LC (390 ppm CO_2_) for 5 weeks. For representative photographs see Fig. 5. Values are means ± SD (n = 7). Asterisks indicate significant changes to the wild type (\**p* < 0.05, \*\**p* < 0.01), bold letters to the *gldt1-1* single mutant and rhombs to the TRX double mutants in the *gldt1-1* background based on students *t*-test (^#^*p* < 0.05).

| Parameter (HC) |  |  |  |
| --- | --- | --- | --- |
| Genotype | Diameter | Total leaf-count | $F_v/F_m$ |
| Col.0 | 11.23 ± 0.64 | 15.29 ± 1.11 | 0.752 ± 0.004 |
| *trxo1* | 11.14 ± 0.63 | 14.71 ± 1.11 | 0.750 ± 0.007 |
| *trxh2* | 11.17 ± 0.57 | 14.57 ± 1.51 | 0.757 ± 0.005 |
| *gldt1* | 10.63 ± 0.48 | 14.86 ± 0.90 | 0.755 ± 0.009 |
| *adg1* | 5.94 ± 0.32** | 11.43 ± 1.27** | 0.716 ± 0.013** |
| *trxo1xgldt1* | 10.74 ± 0.68 | 14.43 ± 1.72 | 0.749 ± 0.007 |
| *trxh2xgldt1* | 11.31 ± 0.86 | 14.71 ± 1.60 | 0.752 ± 0.010 |
| *triple* | 10.80 ± 0.69 | 14.14 ± 1.35 | 0.753 ± 0.005 |
| *trxo1h2* | 10.49 ± 0.83 | 12.67 ± 1.86 | 0.750 ± 0.006 |
| Parameter (LC) |  |  |  |
| Col.0 | 7.60 ± 0.37 | 12.00 ± 0.82 | 0.744 ± 0.005 |
| *trxo1* | 7.44 ± 0.82 | 12.00 ± 1.00 | 0.742 ± 0.005 |
| *trxh2* | 7.60 ± 0.48 | 11.86 ± 1.68 | 0.744 ± 0.006 |
| *gldt1* | 5.17 ± 0.66** | 12.14 ± 1.07 | 0.722 ± 0.007** |
| *adg1* | 4.50 ± 0.59** | 9.57 ± 0.79** | 0.732 ± 0.005** |
| *trxo1xgldt1* | **4.11 ± 0.41\*\*** | **8.57 ± 0.79\*\*** | **0.710 ± 0.009\*\*** |
| *trxh2xgldt1* | **3.16 ± 0.86\*\*** | **9.86 ± 1.35\*\*** | **0.681 ± 0.028\*\*** |
| *triple* | **2.40 ± 0.14\*\*<sup>#</sup>** | **8.17 ± 0.75\*\*** | **0.663 ± 0.025\*\*** |
| *trxo1h2* | 6.80 ± 0.46** | 10.71 ± 1.11** | 0.746 ± 0.003 |

### The impact of multiple *TRX*/*GDCT* mutations on photorespiratory GDC

Targeted metabolite quantification of mtLPD1-dependet pathways was next performed through LC-MS analysis in the full mutant set. To distinguish between metabolic consequences resulting either from impaired regulation of photorespiration or deregulated TCA-cycle, and between short- and long-term acclimation effects, plant material at growth stage 5.1 (Boyes et al., 2001) was analyzed under three different conditions: **(i)** 9 h illumination (end of day - EoD) in HC, **(ii)** 9 h illumination (EoD) with active photorespiration following a shift to ambient air (LC-shift) and **(iii)** 9 h illumination (EoD) with active photorespiration following continuous growth in LC.

Primarily, we aimed to gain insights on *in planta* GDC performance. In HC, the single and double *TRX* mutants displayed largely unaltered glycine and serine amounts, whilst *gldt1* showed a 2-fold increase in glycine and a slight elevation in serine. However, *trxo1h2* and the triple mutant showed a further, but compared to strong photorespiratory mutants (e.g. Timm et al., 2012; Eisenhut et al., 2017) still moderate, elevation in glycine (significant in the triple mutant) under HC conditions. Serine remained almost invariant except for a significant increase in the triple mutant comparable to that of *gldt1* (Fig. 6A1, B1). Upon a shift from HC to LC (i.e. short-term acclimated to ambient air), all mutants accumulated more glycine. The increase was moderate, but similar among the single and double *TRX* mutants and expectedly highest in *gldt1*. Glycine accumulated to higher levels in both *TRX/GLDT* double mutants compared to *gldt1* and tended to be higher also in the triple mutant. Whilst serine was mostly unchanged in *trxo1*, *trxh2* and *trxo1h2*, it followed the opposite trend as glycine in the other mutants. Decreased serine amounts were observed in the single *gldt1*, the double and triple *TRX*/*GDCT* mutants compared to the wild type. Notably, the drop was significantly stronger in *trxo1*x*gldt1* and *trxh2*x*gldt1* compared to *gldt1 a*nd even pronounced in the triple mutant in comparison with both double mutants (Fig. 6A2, B2). In leaves harvested from plants continuously grown in LC (i.e. long-term acclimated to ambient air) a more diverse picture appeared. The lines *trxh2* and *trxo1h2* were significantly decreased in glycine, whereas *trxo1* showed wild-type-like amounts (Fig 6 A3). By contrast, *gldt1*, *trxo1xgldt1 and trxh2xgldt1* (significantly higher as in *gldt1*) and the corresponding triple mutant (significantly higher compared to both *TRX*/*GDCT* double mutants) displayed strongly increased glycine amounts. In contrast, following a transfer in LC minor alterations in serine (decrease in *trxo1*, elevated in *trxo1h2*) became visible in plants grown in normal air (Fig. 6 B3).

**Figure 6.**
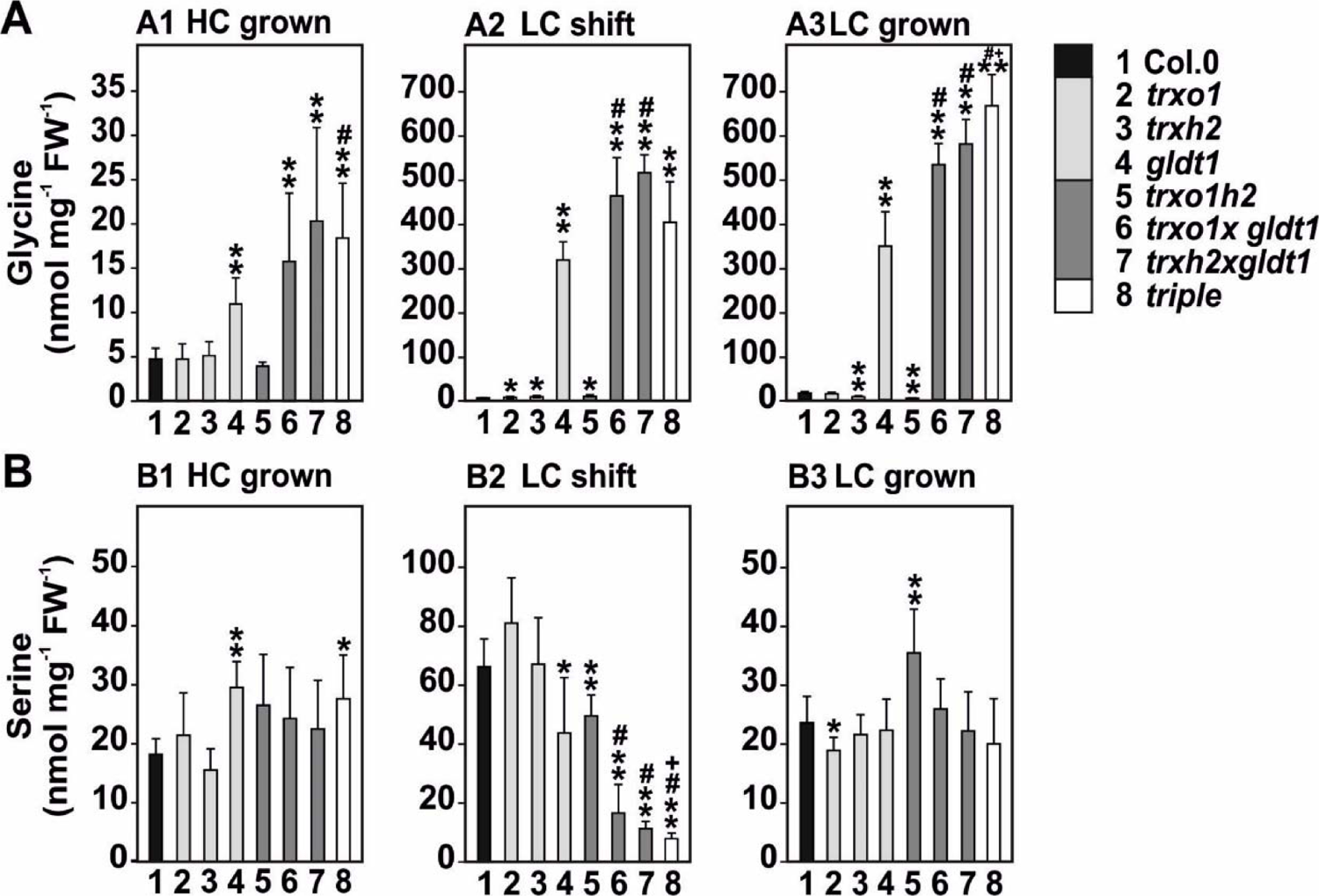
Glycine and serine in the mutant set under different conditions. Absolute **(A)** glycine and **(B)** serine amounts (nmol mg^-1^ FW^-1^) were quantified by LC-MS/MS analysis. Leaf-material was harvested after 9 h of illumination from plants (A1, B1) grown for 8 weeks (growth stage 5.1) under high CO_2_ (HC, 3000 ppm), from plants (A2, B2) shifted from HC to low CO_2_ (LC, 390 ppm; normal air) and (A3, B3) from plants continuously grown in LC. Shown are mean values ± SD (n = 5). Asterisks indicate significant alterations of the mutants compared with the wild type in each condition, rhombs of double *trx*x*gldt1* mutants compared with *gldt1* and plusses of the triple mutant compared to the double *trx*x*gldt* mutants according to Student’s *t*-test (\**p* < 0.05, \*\**p* < 0.01, ^#^*p* < 0.05, ^+^*p* < 0.05).

### Impact of multiple *TRX*/*GDCT* mutations on TCA cycle and BCAAs degradation

Next, we compared metabolic changes in photorespiration intermediates with both other pathways relying on the operation of mtLPD1 containing multienzymes, i.e. the TCA-cycle (PDH, OGDC) and BCAAs degradation (BCKDC). In HC, some TCA-cycle intermediates (citrate, aconitate and 2-oxoglutarate) were decreased in *trxo1*, with minor changes in *trxh2* (decreased aconitate). By contrast, succinate and fumarate were significantly increased in *trxo1*, with no changes in malate and the associated metabolite GABA. In *gldt1* we observed a decrease exclusively in 2-oxoglutarate (Supp. Tab. S1). Among the BCAAs we found only minor changes such as higher valine and isoleucine in *gldt1*. In the multiple mutants, *trxo1h2* showed the fewest changes including a decrease in 2-oxoglutarate and elevated leucine. All lines combining a *TRX* mutation with the *GLDT* knockdown showed the strongest, but not always consistent changes. In *trxo1xgldt1* we found an increase in the TCA-cycle intermediates pyruvate, succinate and fumarate. The *trxh2xgldt1* mutant showed a different response given only citrate was significantly lower. Finally, and in general, the metabolite alterations in the triple mutant in HC were almost comparable to that of *trxo1xgldt1* (Supp. Tab. S1).

Following induction of photorespiration upon a shift from HC-to-LC, the metabolic response of the single mutants regarding the TCA-cycle differed from that observed in HC. After 9 h in air, we observed increased citrate and aconitate in *trxo1* and decreased pyruvate and citrate in *trxh2*. No significant changes occurred in either mutant in the levels of BCAAs. *Gldt1* showed a decrease in pyruvate and citrate and elevated 2-oxoglutarate and succinate. Similar to previous observations, impairment of the photorespiratory flux in *gldt1* caused accumulation of BCAAs (Supp. Tab. S2). Among the multiple mutants, the most interesting change was increased pyruvate in *trxo1h2* after HC-to-LC shift. Most of the other TCA-cycle metabolites (2-oxoglutarate, succinate and GABA) and all BCAA were decreased. The strongest alterations in both pathways were seen in the multiple mutants lacking either one or both *TRX* in the *gldt1* background. However, the responses were comparable between *trxo1xgldt1*, *trxh2xgldt1* and the triple mutant. Hence, all multiple mutants showed accumulation of almost all TCA-cycle intermediates and the BCAAs (Supp. Tab. S2) upon the HC-to-LC shift.

If plants were grown in ambient air (long-term acclimated to photorespiratory conditions), all single mutants showed a few more, partially consistent, changes. With regards to the TCA-cycle, *trxh2* and *gldt1* displayed a similar metabolic signature (increases in most intermediates), whereas *trxo1* was only altered in succinate and malate (higher) and fumarate (lower). However, accumulation of BCAAs was only observed in *trxo1* and *gldt1*, while *trxh2* was similar to the wild type (Supp. Tab. S3). Interestingly, in *trxo1h2* some of the changes seen in the single mutants were not observed anymore. For example, fewer TCA-cycle intermediates were altered (except for pyruvate, succinate, fumarate and malate) and no change occurred in BCAAs. Finally, all multiple mutants in the *gldt1* background displayed a very similar metabolite accumulation pattern. All TCA-cycle intermediates (except fumarate) and BCAAs are present in significantly higher amounts compared to the wild type and, in most cases, also to the single mutants. Nevertheless, the triple mutant showed a few additive effects such as stronger changes in 2-oxoglutarate and fumarate compared to both *TRX*/*GLDT* double mutants (Supp. Tab. S3).

### *In planta* GDC operation following short-term induction of photosynthesis

Our previous studies and the present work provided evidence that the absence of TRX *o1* and *h2* resulted in increased cellular glycine levels under certain environmental conditions, while biochemical analysis indicated that TRX directly regulates (inhibit) the enzymatic activation state of mtLPD1 *in vitro* (Reinholdt et al., 2019a; daFonseica-Pereira et al., 2020). The latter finding suggest that absences of both TRX proteins eventually facilitates mtLPD operation on a short-term and the glycine increase at later stages is due to an inhibition of overall GDC activity. In order to analyze whether increased metabolite pools originate from higher synthesis or slower consumption, a ^13^C-isotope tracing experiment was performed to specifically profile the GDC reaction on a short-term. We restrict our analysis on mtLPD1-depending GDC, because the photorespiratory flux accounts for the major carbon flux in mitochondria upon illumination (e.g. Bauwe et al. 2012). First, the wild type, *gldt1*, the *trxo1h2* double and the corresponding triple mutant plants were grown in HC for 5 weeks. Subsequently, plant leaves were fed with ^13^C-glycine in HC and LC to distinguish between conditions with suppressed or active photorespiration. The ^13^C-glycine feeding was done for 30 and 60 min under illumination. Metabolites were isolated from frozen leaves and the incorporation of ^13^C in glycine and serine followed by gas chromatography coupled to mass spectrometry (GC-MS) analysis.

Under control conditions (dark, HC), we detected no significant differences in the natural ^13^C-glycine pattern among all genotypes. As expected, the fractional ^13^C-enrichment following illumination, i.e. induction of photosynthesis, in glycine is significantly lower in all genotypes in HC compared to LC conditions (Fig. 7A). However, *trxo1h2* displayed slightly lower (significant after 60 min) ^13^C-enrichments in glycine, whereas *gldt1* (significant after 30 and 60 min) and the triple (significant after 30 min) mutant showed higher ^13^C-glycine accumulation after onset of illumination at HC than wild type (Fig. 7A). The effects seen in HC became much more distinct with plants material incubated under LC conditions. LC-exposed wild type displayed an almost linear, much higher, increase in the ^13^C-enrichment in glycine compared to HC. Interestingly, *trxo1h2* also showed an increase in ^13^C in glycine but it was about half as much as observed in the wild type. Finally, *gldt1* and the triple mutant showed significantly elevated amounts of ^13^C-glycine compared to the control which culminated in an about 5-times higher fraction after 60 min in both lines grown compared to wild type under LC conditions (Fig. 7A). If one considers serine, the product of the GDC and serine-hydroxymethyltransferase reaction, ^13^C-enrichment in the course of the experiment was also observed, but the enrichment rate was much lower compared to glycine (Fig. 7B). Moreover, less strong changes were seen in both conditions and among the genotypes after 60 min of ^13^C-glycine feeding, except a significant increase in the ^13^C-enrichment in serine in *trxo1h2* after 60 min (Fig. 7B). These findings suggest that GDC activity is increased in the *trxo1h2* double mutant (less glycine and enhanced serine label) on a short-term, whilst as expected it is already decreased under these conditions in the *gldt1* background.

**Figure 7.**
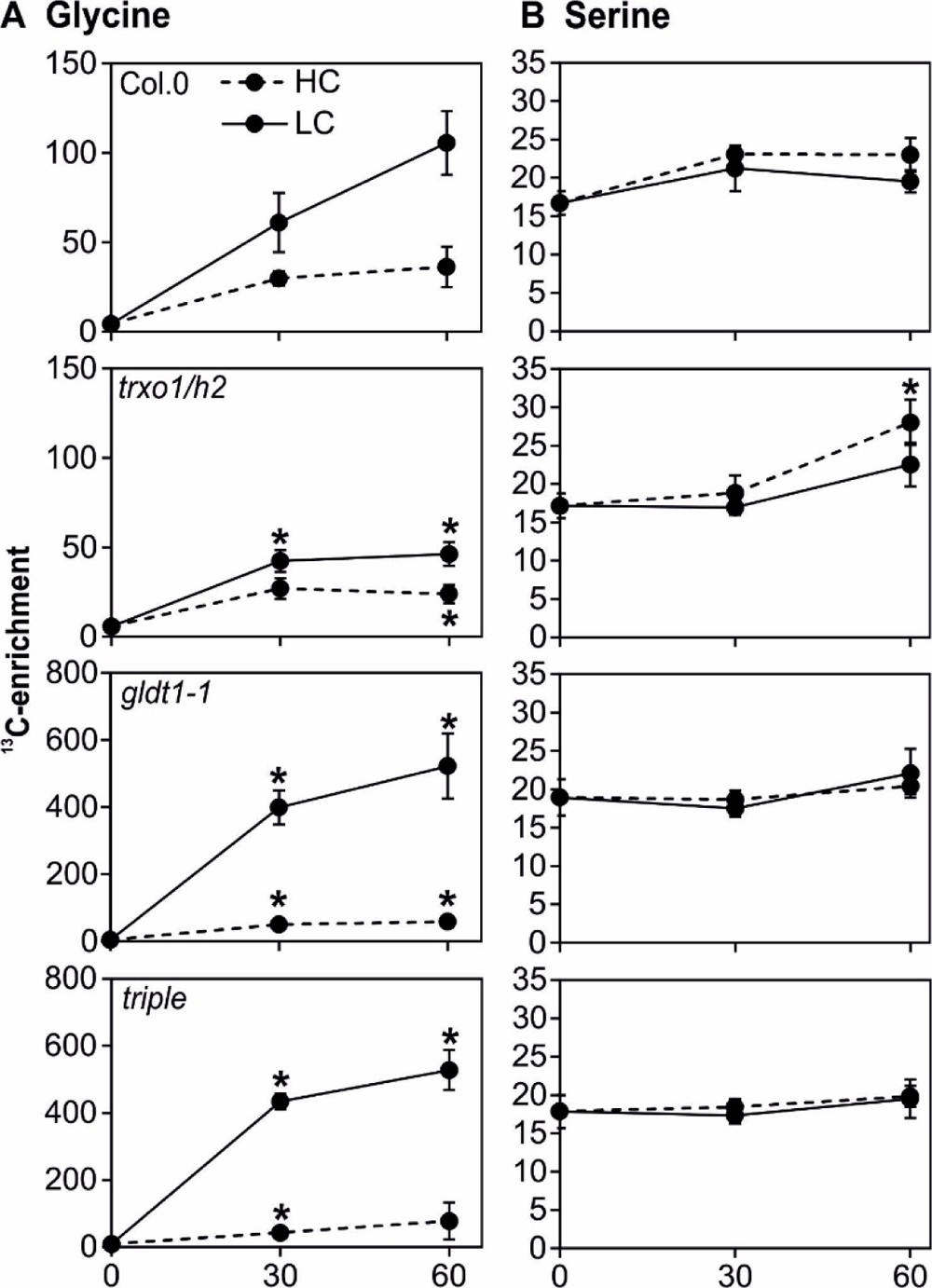
^13^C-enrichment in glycine and serine in selected mutants in HC and LC. Before onset of illumination, plant leaf’s were cut from the rosettes in the dark following 5 weeks of growth in HC conditions. Subsequently, leaf’s were fed with ^13^C-glycine through the petiole for 30 and 60 min of light in HC (dashed line) and LC (solid line) conditions. ^13^C-enrichment in **(A)** glycine and **(B)** serine was followed through GC-MS analysis. Shown are mean values ± SE (N > 4). Asterisks indicate significant alterations of the mutants compared with the wild type in each condition according to Student’s *t*-test (\**p* < 0.05). For further experimental details see material and methods section.

### Pyridine nucleotide contents in multiple *TRX*/*GDCT* mutants under different conditions

Regulation of redox-reactions in primary metabolism through the cellular TRX system mainly responds to the redox state of the cell, i.e. to changes in the relative reduction levels of pyridine nucleotides. Therefore, we next quantified the amounts of NAD(H)^+^ and NADP(H)^+^ in the mutants under the same conditions (HC, shifted to LC and grown in LC) and time points (EoD) used for the metabolite analyses above. In HC, we detected no significant changes in NADH, NAD^+^ or the respective NADH/NAD^+^ ratios among the genotypes (Table 2, Supp. Tab. S4), which is consistent with the lower impact of *TRX* mutations on metabolism under this condition. Despite NADPH was largely unaltered in all mutants, NADP^+^ was significantly decreased in *trxh2*, *gldt1*, *trxo1h2*, *trxh2xgldt1* and the triple mutant. However, these changes translated to significantly altered NADPH/NADP^+^ ratios only in *gldt1* and *trxo1h2* (Table 2, Supp. Tab. S4). Upon the shift to LC, *trxo1xgldt1* was found to be increased in NADH whilst the NADH/NAD^+^ ratio was higher in *trxh2*, *trxo1xgldt1* and the triple mutant compared to the wild type. With regards to the phosphorylated pyridine nucleotides only few changes were seen. This included *trxo1* among the single mutants (decreased NADPH and NADPH/NADP^+^ ratio) and the *trxo1h2* double mutant (decreased NADPH and NADP^+^). All other mutants displayed values comparable to the wild type (Table 2, Supp. Tab. S5). Finally, if leaf-material was analyzed from plants continuously grown in LC, more systematic changes emerged. Despite NAD^+^ was only decreased in *trxo1*, almost all other mutants (except *trxh2* and *trxo1xgldt1*) contained significantly lower amounts of NADH. These changes caused a noticeable reduction in the NADH/NAD^+^ ratio in almost all of the analyzed genotypes, excluding *trxh2* (Table 2, Supp. Tab. S6). At one hand, NADPH tended to be increased in some of the multiple mutants, however, these changes were not significant compared to the control. On the other hand, all three single mutants and the *trxh2xgldt1* and the *trxo1h2* double mutants displayed decreased amounts of NADP^+^. All the alterations mentioned before ultimately caused a significant increase in the NADPH/NADP^+^ ratio in *gldt1*, *trxh2xgldt1*, *trxo1h2* and the triple mutant (Table 2, Supp. Tab. S6).

**Table 2.** Pyridine nucleotide ratios in the mutant set under different conditions. Pyrimidine nucleotide contents were determined from plants grown to growth stage 5.1 (Boyes et al., 2001) at the end of the day (EoD, 9 h illumination) in HC, following a shift from HC-to-LC and in LC grown plants. Values are means ± SD (n = 6) and bold letters indicate values statistically significant from the wild type based on Students *t*-test (\**p* < 0.05, \*\**p* < 0.01). For single values please see Supplemental Tables 4-6.

| Genotype | NADH/NAD<sup>+</sup> | NADH/NAD<sup>+</sup> | NADH/NAD<sup>+</sup> |
| --- | --- | --- | --- |
|  | HC | LC-shift | LC |
| Col.0 | 0.119 ± 0.008 | 0.072 ± 0.018 | 0.136 ± 0.017 |
| *trxo1* | 0.144 ± 0.027 | 0.074 ± 0.029 | 0.108 ± 0.024 |
| *trxh2* | 0.139 ± 0.031 | **0.103 ± 0.012\*** | 0.116 ± 0.039 |
| *gldt1* | 0.136 ± 0.025 | 0.086 ± 0.021 | 0.108 ± 0.011 |
| *trxo1xgldt1* | 0.118 ± 0.020 | **0.114 ± 0.018\*** | **0.099 ± 0.001\*\*** |
| *trxh2xgldt1* | 0.128 ± 0.020 | 0.089 ± 0.016 | **0.102 ± 0.010\*\*** |
| *triple* | 0.130 ± 0.017 | **0.112 ± 0.015\*** | **0.096 ± 0.015\*** |
| *trxo1h2* | **0.145 ± 0.010\*\*** | **0.106 ± 0.016\*** | **0.108 ± 0.017\*** |
|  | NADPH/NADP<sup>+</sup> | NADPH/NADP<sup>+</sup> | NADPH/NADP<sup>+</sup> |
|  | HC | LC-shift | LC |
| Col.0 | 0.052 ± 0.013 | 0.609 ± 0.121 | 0.077 ± 0.011 |
| *trxo1* | 0.047 ± 0.016 | **0.460 ± 0.082\*** | 0.081 ± 0.018 |
| *trxh2* | 0.057 ± 0.013 | **0.423 ± 0.058\*\*** | 0.080 ± 0.023 |
| *gldt1* | 0.067 ± 0.008 | 0.617 ± 0.082 | **0.101 ± 0.018\*** |
| *trxo1xgldt1* | 0.052 ± 0.007 | 0.678 ± 0.118 | **0.109 ± 0.015\*\*** |
| *trxh2xgldt1* | 0.060 ± 0.010 | 0.650 ± 0.088 | **0.129 ± 0.011\*\*** |
| *triple* | 0.057 ± 0.015 | 0.697 ± 0.096 | **0.104 ± 0.016\*** |
| *trxo1h2* | 0.072 ± 0.015 | 0.656 ± 0.112 | **0.100 ± 0.017\*** |

Taken together, the absence of different TRX proteins had marked impact on the relative reduction of pyridine nucleotides. Especially, the response of the NADH/NAD^+^ ratio is biphasic given it is mostly increasd after short-term, but decreased, after long-term acclimation to photorespiratory conditions in most mutants.

### Plant mtLPD1 is inhibited by NADH

TRX *o1* and *h2*, respectively, were shown to contribute to the regulation of the activation state of mtLPD1 *in vitro* and, perhaps, *in vivo* (Reinholdt et al., 2019a; daFonseica-Pereira et al., 2020). However, since TRX *h2* was reported to reside to the microsomal fraction rather than to mitochondria (Hou et al., 2021), the impact of TRX *h2* deficiency on mtLPD1 is likely indirect, presumably via modulation of the mitochondrial redox state. Interestingly, a study of *E. coli* LPD revealed its inhibition by NADH at physiological concentrations (Kim et al., 2008), making this molecule a prime candidate for further regulation of mtLPD1 in response to altered mitochondrial pathway fluxes, especially photorespiration. For this reason, and the changes observed in the NADH/NAD^+^ ratio in the mutants (Table 2), recombinant mtLPD1 was obtained from *Pisum sativum* (*Ps*mtLPD1) through heterologous expression and affinity purification and the activity tested with different NADH concentrations as potential inhibitor. The biochemical analysis revealed a considerable inhibition of *Ps*mtLPD1 by NADH (Fig. 8), whilst NADPH had no effect (data not shown). The inhibitor constant (*K_i_*) for NADH estimated from these measurements was ∼ 66.11 µM, which is comparable to the one recently reported for the recombinant LPD from the cyanobacterium *Synechocystis* sp. PCC 6803 (Wang et al., 2022).

**Figure 8.**
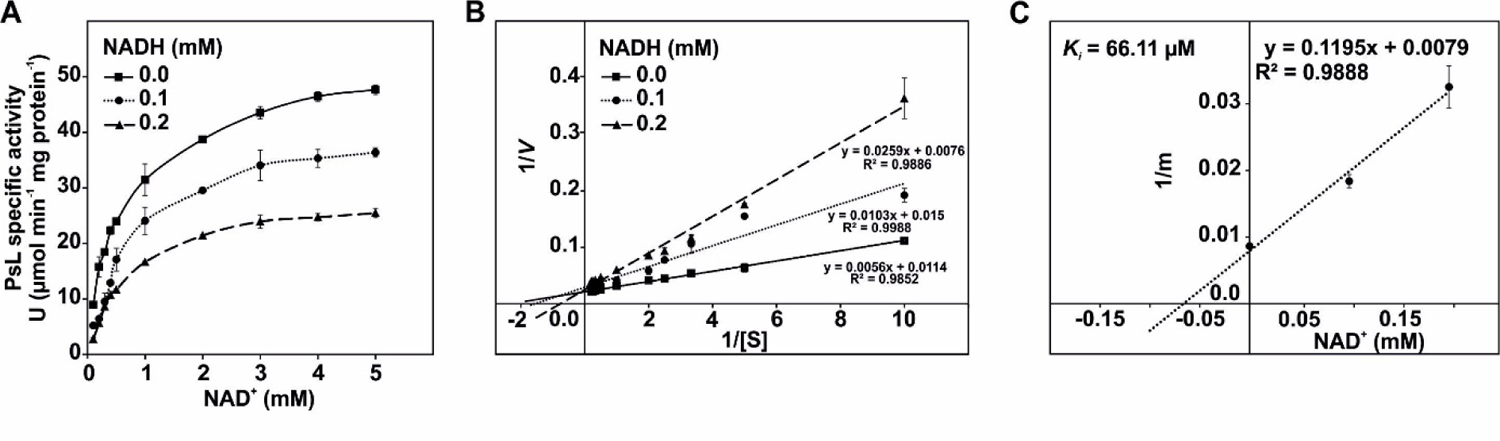
Inhibition of *Pisum sativum* mtLPD1 (PsL) by NADH. **(A)** The rate of PsL activity was measured in the forward direction (3 mM DL-dihydrolipoic acid) as a function of NAD^+^ (0.1, 0.2, 0.3, 0.4, 0.5, 1, 2, 3, 4 and 5 mM) with the indicated NADH concentrations (0, 0.1 and 0.2 mM). Specific enzyme activity is expressed in µmol NADH per min^-1^ mg protein^-1^ at 25°C. **(B)** Lineweaver-Burk plots of the three NADH concentrations. **(C)** The inhibitor constant (*K_i_*) was estimated by linear regression of (1) the slopes of the three Lineweaver-Burk plots at the three NADH concentrations versus (2) the NADH concentration. Shown are mean values ± SD from at least three technical replicates.

## Discussion

Redox regulation through TRXs is a fundamental mechanism to regulate a plethora of biological processes including metabolic fluxes, stress acclimation and tolerance or protein transport and processing (Balmer et al., 2004; Buchanan, 2016; Geigenberger et al., 2017; Møller et al., 2020). Next to their canonical chloroplastidal functions, recent research revealed TRXs also impact on mitochondrial processes. Hence, TRX *o1* and *h2* affect the operation of the TCA-cycle, photorespiration and, perhaps, the degradation of BCAAs (Daloso et al., 2015; Reinholdt et al., 2019; Møller et al., 2020; Timm and Hagemann, 2020; daFonseica-Pereira et al., 2020; 2021). Our main goal here was to test if TRX *o1* and *h2* mediated regulation of the three mitochondrial pathways could be explained through a regulatory mechanism involving a shared enzymatic activity/protein, namely mitochondrial dihydrolipoamide dehydrogenase (mtLPD1).

### On the similarities and differences between the effects of deficiencies in TRX o1 and h2

First, we aimed to find out if TRX *o1* and *h2* share a common or a specific response among the afore mentioned mtLPD1-dependet mitochondrial pathways. If they share a common response, we would expect similar metabolite patterns with both *TRX* single mutants. To prove this hypothesis, we designed experiments to explicitly induce short-term environmental fluctuations through triggering the necessity for optimal photorespiration (Fig. 1A). This was mainly performed since photorespiration represents the main mitochondrial flux in illuminated leaves (Bauwe et al., 2012; Lim et al., 2020) and deregulation of mtLPD1 might be best visible under these conditions. Indeed, we found a rather consistent metabolite accumulation pattern in the *trxo1* and *trxh2* single mutants among representative intermediates of photorespiration and BCAAs degradation. The pattern correlates with the rate of photorespiration since no significant change was seen in darkness or elevated CO_2_ (Fig. 1B, C). The statement is in close agreement with the observed changes in levels of TCA-cycle intermediates since most metabolite patterns were comparable among the single mutants, with some specific variations (Fig. 2). Such discrepancies are, however, best explained by the different subcellular localization of both TRXs and the specific enzyme targets of TRX *o1* such as SDH and fumarase (Daloso et al., 2015; Hou et al., 2021). The direct impact on both enzymes was discussed to support the regulation of the different flux modes of the TCA-cycle and its inactivation in the light in addition to PDH regulation (Sweetlove et al., 2010; Araújo et al., 2011; Daloso et al., 2015; Lima et al., 2021). Interestingly, the latter statement is further supported by our metabolite analysis since pyruvate significantly accumulates in both TRX single mutants on a short-term (Fig. 2). Hence, these experiments augment our supposition that TRX *o1* and *h2* might signal through a similar regulatory mechanism.

### Are there additive effects through lack of TRX o1 and h2 on mitochondrial functions?

The next question we tackled was if combined deficiency of TRX *o1* and *h2* has additive effects on mitochondrial functions. This was of interest because of the high redundancy in the regulation circuits involving TRX and glutaredoxins (Reichheldt et al., 2005). Indeed, the double *trxo1h2* mutant (Hou et al., 2021) showed significantly stronger impairment of the photosynthetic-photorespiratory gas exchange compared to single mutants (Fig. 3; Reinholdt et al., 2019; daFonseica-Pereira et al., 2020). Especially the increase in γ, a common classic diagnostic parameter for impaired photorespiration, was significantly elevated (∼10% higher) compared to the single deletions. The responses were comparable to studies with other intermediate photorespiratory mutants (Timm et al., 2011; Cousins et al., 2011; Timm and Bauwe, 2013) and, thus, corroborated the photorespiration defect. Another line of evidence arose from the growth and metabolite analysis following short-term environmental perturbation. The *trxo1h2* double mutant showed clear signs of increased O_2_-sensitivity on the phenotypic level (Fig. 5B; Supp. Fig. 1) and a stronger slowdown of the photorespiratory metabolite conversion (Fig. 4). Compared to the single mutants that also accumulated glycine (Fig. 1B), deletion of both *TRX* significantly exacerbated these changes (Fig. 4). Interestingly, the GDC impairment did not resulted in major changes in serine, suggesting that a potential depletion can be prevented on the short-term potentially via the operation of other serine biosynthetic routes (Ros et al., 2014). Collectively, these results corroborated additive impairment of photorespiration in *trxo1h2*, presumably GDC. This assumption could be further supplemented with mutants exhibiting boosted GDC impairment through single and double TRX deficiency in the *gldt1* mutant background (Timm et al., 2018). In comparison with the *trxo1xgldt1* double mutant analyzed before (Reinholdt et al., 2019a), growth of *trxh2xgldt1* was similarly impaired. Notably, simultaneous deletion of TRX *o1* and *h2* in *gldt1* (triple mutant) caused significantly stronger symptoms (Table 1, Fig. 5A). However, all effects were mainly caused through a photorespiration defect given most alterations normalized in HC-grown plants, suppressing photorespiration (Table 1, Fig. 5B).

### Metabolic responses of mtLPD1-dependent pathways in multiple mutants

The above results pointed to additive phenotypic and physiological responses through lack of both *TRXs* in the wild type and the *gldt1* mutant. Therefore, we expected a coincided impact also on mtLPD1-dependet pathways. Targeted metabolomics, revealed this holds true mostly for photorespiration and BCAAs degradation, but responses differ if plants were acclimated to ambient air on a short- or long-term. Similar to the growth effects, minor metabolic changes were seen in HC (Fig. 6, Supp. Table 1), which agrees with previous studies on *gldt1* and *trxo1xgldt1* (Timm et al., 2018; Reinholdt et al., 2019). Impaired photorespiratory GDC (glycine accumulation) largely mirrored the strength of the mutation in response to short-term ambient air acclimation (minor increases in the single mutants and *trxo1h2* < intermediate in *gldt1* < highest in *TRX* and *GLDT* double and triple mutants). However, the increases were rather similar among the multiple genotypes involving *TRX* and *GLDT1* mutations and additive effects in the triple mutant were only seen in serine on a short-term (Fig. 6A, 6B) or in glycine if plants were long-term acclimated to photorespiratory conditions (Fig. 6C). Serine alterations largely normalized if plants were grown in ambient air, again suggesting other serine biosynthesis pathways can compensate for a lowered GDC turnover (Ros et al., 2014). Similar to glycine, a general and comparable increase of all three BCAAs was seen in all genotypes harboring the *gldt1* mutation. The accumulation was strongest upon the HC-to-LC shift (short-term induction of photorespiration), including some additive effects in mutants with one or both TRX in *gldt1* compared to the *gldt1* (Supp. Table 2, 3). Impaired BCAAs degradation is shared with classic photorespiration mutants (Timm et al., 2012) and once more serves as evidence for impaired photorespiration and a metabolic connection of both processes mentioned earlier (Florian et al., 2013; Dellero et al., 2021). Interestingly, a clear variation of this response was a consistent decrease of all amino acids in *trxo1h2* after 9 h in air, but not in air grown plants, suggesting faster turnover on a short-term (Supp. Table 2, 3).

TCA-cycle intermediates did not always respond in a consistent manner across the mutants in comparison with photorespiration and BCAAs degradation. However, this finding is not surprising considering the different possible flux-modes (Sweetlove et al., 2014) and multiple regulation sites (Nunes-Nesi et al., 2013; Zhang et al., 2021), in the pathway. In addition to the mtLPD1-dependent enzyme complexes, namely PDH and OGDC (Millar et al., 1998; 1999), TRX *o1* can directly act on SDH and fumarase (deactivating) and the associated enzyme ATP-citrate lyase (ACL, activating) (Daloso et al., 2015). During illumination, and in HC, we expect either, very low rates of photorespiration and restricted carbon influx into the TCA-cycle because of inhibited PDH through phosphorylation (Tcherkez et al., 2005; Zhang et al., 2021). Consequently, the impact of impaired photorespiration and the subsequent malfunctioning of mtLDP1-dependet regulation of enzyme complexes should be lower. In this regard, pyruvate accumulated only in the stronger mutants (*trxo1xgldt1-1* and triple). 2-Oxoglutarate was decreased in several genotypes (*trxo1*, *gldt1*, *trxo1xh2*, triple) which, perhaps, might be indicative for decreased flux through nitrogen assimilation. The clearest response on the TCA-cycle in HC was in *trxo1*, since intermediates of the oxidative branch (citrate, aconitate and 2-oxoglutarate) were lower, whilst some of the reductive branch (succinate and fumarate) tended to accumulate (Suppl. Table S1). Such behavior is congruent with the specific enzyme targets (Daloso et al., 2015). In conditions with active photorespiration, and in terms of a uniform response, all mutants with either one or two deleted *TRX* in the *gldt1* background displayed strongly impaired operation of the TCA-cycle. This is already seen after the LC-shift but was most distinct in air grown plants (Suppl. Table 2, 3). Due to the specific involvement of mtLPD1 in PDH and OGDC, we mostly considered alterations in their substrates. Indeed, *gldt1* and most multiple mutants (except *trxo1h2*) showed the strongest responses in pyruvate and especially in 2-oxoglutarate. This finding suggests that either, OGDC represents a central regulatory target or that nitrogen metabolism (i.e. ammonia refixation) is also affected through deletion of *TRX* and *GLDT*. The latter statement is congruent with many studies on photorespiratory mutants, either through perturbation of the photorespiratory nitrogen cycle and nitrogen metabolism in general (Bloom et al., 2010; Bloom, 2015; Timm et al., 2012; Dellero et al., 2015). Further work is needed in order to comprehensively characterize nitrogen metabolism in *TRX* deficient mutants with regards to elucidating the underlying regulatory mechanisms.

### Redox regulation of mtLPD1 via TRX o1 and h2 allows a concerted regulation of multiple mitochondrial pathways

In terms of a consistent regulatory mechanism, our prime target for simultaneous effects on photorespiration, the TCA-cycle and BCAAs degradation was mtLPD1, as a shared enzyme (Bourguignon et al., 1988; Millar et al., 1998; 1999). This assumption originated from the previous finding that TRX *o1* and *h2* regulate mtLPD1 *in vitro* (Reinholdt et al., 2019; daFonseica-Pereira et al., 2020) and the additive negative effects on the double mutant observed during this study (e.g. Fig. 3-6). However, since only TRX *o1* localizes to the mitochondria (Laloi et al., 2001) and TRX *h2* resides to the microsomal fraction under normal conditions (Hou et al., 2021; Lee et al., 2021), direct regulation of the enzymatic activity can only occur through TRX *o1*. The central question then was how to connect the different compartments with an underlying regulatory mechanism affecting multiple mitochondrial pathways. As another possibility we assumed parallel regulatory impact on the three pathways through mtLPD1 via a metabolic signal in response to changes in the redox states of subcellular compartments involving TRX *o1* and *h2*. The latter hypothesis is congruent with alterations in the pyrimidine nucleotide amounts observed previously (Reinholdt et al., 2019; daFonseica-Pereira et al., 2020) and the recent finding of considerable shifts in the redox couples of ascorbate and glutathione in the *TRX* mutants, especially *trxo1/h2* (Calderón et al., 2018; Hou et al., 2021). During this study we mainly found consistent biphasic changes in the NADH/NAD^+^ ratio, i.e. significant increases after 9 h in ambient air and significant decreases in plants continuously grown in ambient air (Table 2, Suppl. Tables 4-6). This result might be indicative for a short-term overproduction of NADH through increased mtLPD1 activity, which became attenuated after long-term acclimation to photorespiratory conditions. This statement agrees well with the glycine accumulation under the different conditions (Fig. 1B, 4, 6 and 7A). Changes in the NADPH/NADP^+^ ratio, which is more representative for chloroplastidal metabolism, were not so systematic and mainly restricted to mutants with the *gldt1* background. The most likely explanation for these changes is a slowdown of photosynthesis, i.e. the Calvin-Benson cycle, due to impaired photorespiratory metabolism and the resulting feedback inhibition through pathway intermediates. However, in light of the changes in the cellular NADH/NAD^+^ ratios (Table 2), and based on two previous studies showing inhibition of LPD from *E. coli* (Kim et al., 2008) and Synechocystis (Wang et al., 2022), we tested plant mtLPD1 towards its susceptibility for NADH. Pea mtLPD1 was taken since it was already available from previous work (Reinholdt et al., 2019a) and we failed to obtain soluble mtLPD1 from Arabidopsis. As shown in Fig. 8, PsmtLPD1 was inhibited by NADH at physiological concentrations, too. The estimated *K_i_* was ∼66.11 µM – being similar in range to those reported for the *E. coli* and Synechocystis enzymes (Kim et al., 2008; Wang et al., 2022). The biphasic consequences of impaired TRX operation is also supported through the ^13^C-labeling approach to profile *in planta* GDC activity following 30-to-60 minutes of illumination (Fig. 8). The *trxo1h2* double mutant was compared with *gldt1* in order to differentiate between effects mediated through redox changes with those directly occurring due to the impairment of photorespiration. Interestingly, *trxo1h2* was characterized by decreased fractional ^13^C-enrichment into glycine compared to the wildtype, whilst *gldt1* and the triple mutant showed the anticipated elevated incorporation of the isotope (Fig. 8). From this comparison it seems reasonable to conclude that absence of both TRXs facilitates mtLPD1 and, in turn, overall GDC activity in the very short-term, leading to a massive overproduction of NADH after induction of photorespiration. Consequently, the mitochondrion becomes more reduced and the NADH/NAD^+^ ratio rises. Interestingly, such a scenario would ultimately inhibit GDC activity as reported previously (Bourguignon et al., 1988) and is likely to explain the high glycine accumulation seen in the multiple mutants (Fig. 6). Finally, this result, in conjunction with our findings presented here, would suggest the mtLPD1 regulation would not only have implication for photorespiratory GDC but also towards all other mtLDP1-dependent pathways.

## Material and methods

### Plant material and growth

In this study, *Arabidopsis thaliana* L. (Arabidopsis) ecotype Columbia.0 (Col.0) served as wild type reference. We used the following, previously isolated and characterized T-DNA insertional lines: *trxo1-1* (SALK 042792, Daloso et al., 2015), *trxh2-2* (SALK-079516, Laloi et al., 2004), the corresponding double mutant *trxo1h2* (Hou et al., 2021), *gldt1-1* (WiscDSLox 366A11-085, Timm et al., 2018) and the *trxo1-1*x*gldt1-1* double mutant (Reinholdt et al., 2019a). Prior sowing on a mixture of soil and vermiculite (4:1), seeds were surface-sterilized with chloric acid. Subsequently, pots were incubated at 4°C for at least 2 days to break dormancy. All plants were grown under controlled environmental conditions as follows: 10 h day, 20°C/ 14 h night, 18°C, ∼120 µmol photons m^-2^ s^-1^ light intensity, 70% relative humidity and with two different CO_2_ concentrations (HC – high CO_2_, 3000 ppm and LC – low CO_2_, 390 ppm). Where specified in the text, plants were shifted from HC-to-LC to study the acclimation to altered CO_2_ concentrations with otherwise equal conditions. Plants were regularly watered and fertilized weekly (0.2% Wuxal, Aglukon).

### Generation and verification of multiple *TRX* and *GLDT* mutants

The *trxo1-1xgldt1-1* T-DNA insertional line was crossed with *trxh2-2* to obtain a *trxh2-2xgldt1-1* double and a *trxo1-1xgldt1-1xtrxh2-2* (triple) mutant. To verify the T-DNA insertions in the corresponding genes, leaf-DNA was isolated according to standard protocols and PCR-amplified (min at 94°C, 1 min at 58°C, 2 min at 72°C; 35 cycles) with primers specific for the left border (R497 [5’-AACGTCCGCAATGTGTTATTAAGTTGTC-3’] for WiscDsLox line, *gldt1-1*, P946 [5’-TGGTTCACGTAGTGGGCCATC-3’] for SALK 042792 line, *trxo1-1*, P959 [5’-ATTTTGCCGATTTCGGAAC-3’] for SALK-079516 line, *trxh2-2*) and *AtGDC-T* (R490 [5’-ACAAAGTCATGGACGAAGGAGACACAC-3’]), *TRX o1* (P945 [5’-CAACACGTTCTTTACTAGACG-3’]) and *TRX h2* (P957 [5’-GATAATGGGAGGAGCTTTATC-3’]) specific primers. The resulting fragments (*gldt1-1* 1859 bp, *trxo1-1* 1200 bp and *trxh2-2* 1100 bp) were sequenced for verification of the T-DNA fragment within the genes. Zygosity was examined by PCR amplification (min at 94°C, 1 min at 58°C, 2 min at 72°C; 35 cycles) of leaf DNA with the primer combination R498 [5’-GTCCCTTTTGCCATTGATAGCAAC-3’] and R490 for *GLD-T* (1943 bp), P944 [5’-CTCGAGTGATGAAGGGAAATT-3’] and P945 for *TRXo1* (1800 bp) and P957 and P958 [5’-CACAAGACTATTGGTTAAGG-3’] for *TRXh2* (1035 bp).

### Gas exchange and chl a fluorescence measurements

Fully expanded rosette leaves of plants at growth stage 5.1 (Boyes et al., 2001) were used to determine leaf gas exchange parameters. Standard measurements were carried out with the following conditions: photon flux density = 500 µmol m^-2^ s^-1^, chamber temperature = 25°C, flow rate = 300 µmol s^-1^, and relative humidity = 60 to 70%. To determine CO_2_ compensation points (Γ), A/C_i_ curves were recorded with the following CO_2_ steps: 400, 300, 200, 100, 50, 0, and 400 ppm. We used the gas-mixing system GMS600 (QCAL Messtechnik) to change the O_2_ concentrations to 3 or 50% (balanced with N_2_). Oxygen inhibition of A was calculated from measurements at 21 and 50% oxygen using the equation: O_2_ inhibition = (A_21_ – A_50_)/A_21_*100. Calculation of γ was performed by linear regression of the Γ versus-oxygen concentration curves and given as slopes of the respective functions. Chlorophyll a fluorescence was determined using a PAM fluorometer (DUAL-PAM-100; Walz). Maximum quantum yields of PSII (*F_v_*/*F_m_*) were determined after transfer into darkness (10 min) of the plants grown under HC and LC conditions.

### Metabolite analysis

The determination of metabolites was carried out by liquid chromatography coupled to tandem mass spectrometry (LC-MS/MS) analysis using the LCMS-8050 system (Shimadzu, Japan). For this purpose, we harvested leaf material (∼25 mg) from fully expanded rosette leaves of plants at growth stage 5.1 (Boyes et al., 2001) grown in HC or LC or following a transition from HC to LC at the end of the day (EoD, 9 h illumination). The material was immediately frozen in liquid nitrogen and stored at −80°C until further analysis. Extraction of soluble primary intermediates was carried out using LC-MS grade chemicals according to the method described in Arrivault et al., (2009; 2015) and the samples analyzed exactly as described in Reinholdt et al. (2019a). The compounds were identified and quantified using multiple reaction monitoring (MRM) according to the values provided in the LC-MS/MS method package and the LabSolutions software package (Shimadzu, Japan). Authentic standard substances (Merck, Germany) at varying concentrations were used for calibration and peak areas normalized to signals of the internal standard ((morpholino)-ethanesulfonic acid (MES), 1 mg/ml). Data were interpreted using the Lab solution software package (Shimadzu, Japan).

### Determination of pyridine nucleotide contents

For these measurements we used plants at growth stage 5.1 (Boyes et al., 2001) and the same time points as stated for the metabolite analysis. The contents of NAD^+^/NADH and NADP^+^/NADPH were determined in acid and alkaline extracts using the protocol described in Zhang et al (2020). The assays involve the phenazine methosulfate-catalysed reduction of dichlorophenolindophenol (DCPIP) in the presence of ethanol and alcohol dehydrogenase (for NAD^+^ and NADH) or glucose-6-P and glucose-6-P dehydrogenase (for NADP^+^ and NADPH). Reduced and oxidized forms are distinguished by preferential destruction through measurements in acid or alkaline buffers.

### ^13^C-labeling and GC-MS analysis

For the ^13^C-isotope labeling approach, we used 5-week-old plants grown under standard conditions defined above but with high CO_2_ (HC – 3000 ppm). Before onset of illumination, intact leaf’s were cut from the plant rosette with a razor blade under weak green light to prevent induction of photosynthesis and photorespiration. Subsequently, the leaf’s were put into Eppendorf tubes containing 10 mM [U-^13^C]-glycine in 10 mM MES-KOH solution, pH 6.5, and placed back into growth cabinets. Following 30- and 60-min illumination in normal air (LC - 400 pmm CO_2_) and HC, the material was harvested after the petiole was cut from the leaf to prevent contamination with residual ^13^C label, flash frozen in liquid nitrogen and stored in −80°C until further analysis. Metabolite extraction and GC-MS analysis were carried out exactly as described earlier (Lisec et al., 2006). Metabolite identification was carried out using the Golm Metabolome Database (http://gmd.mpimp-golm.mpg.de/) (Kopka et al., 2005). The detection of the isotopologues was made by using the software Xcalibur® 2.1 (Thermo Fisher Scientific, Waltham, MA, USA). The fractional ^13^C-enrichment was then performed as described earlier (Lima et al., 2021).

### Expression, purification and activity measurements of recombinant mtLPD1

To obtain recombinant mtLPD1 from *Pisum sativum* for biochemical analysis we followed the same procedure described in Reinholdt et al. (2019a). The activity of mtLPD was assayed spectrophotometrically at 340 nm in the forward direction as described previously (Timm et al., 2015) using 5-10 µg ml^-1^ recombinant protein to initiate the reaction. Enzyme activity was recorded with three different NADH concentrations (0, 100 and 200 µM) and is expressed in µmol NADH per min^-1^ mg protein^-1^ at 25°C.

### Statistical analysis

We used the two-tailed Student’s *t* test (Microsoft Excel 10.0) and ANOVA analysis (Holm and Sidak test; Sigma Plot 11; Systat Software) for multiple genotypes for comparisons. The term significant is used only if the change in question was confirmed to be significant at the level of \**p* < 0.05 and \*\**p* < 0.01.

## Accession numbers

The Arabidopsis Genome Initiative or GenBank/EMBL database contains sequence data from this article under the following accession numbers: *ADG1* (At5g48300), *TRX o1* (At2g35010), *TRX h2* (AT5G39950), *GLDT* (At1g11860) and pea *mtLPD1* (P31023).

## Acknowledgements

We wish to thank Klaudia Michl (Rostock) and Ina Krahnert (Golm) for valuable technical assistance. The gift of the *adg1-1* mutant by Hans-Henning Kunz (München) and *gldt1-1* mutant by Hermann Bauwe (Rostock) is gratefully acknowledged. We thank the Nottingham Arabidopsis Stock Centre for donation of the T-DNA insertional lines. Work in the authors Lab was financially supported by the University of Rostock (S.T. and M.H.). The LC-MS equipment at the University of Rostock was financed through the HBFG program (GZ: INST 264/125-1 FUGG to M.H.). S.A, Y.Z and A.R.F would like to thank the Max-Planck Society and European Union’s Horizon 2020 research and innovation program, project PlantaSYST (SGA-CSA No 664621 and No 739582 under FPA No. 664620) for supporting their research. P.G. and L.-Y.H. gratefully acknowledge support by the Deutsche Forschungsgemeinschaft (TRR 175, The Green Hub). We further thank the research fellowship granted by the National Council for Scientific and Technological Development to DMD (CNPq, 303709/2020-0) and the scholarship granted by the Brazilian Federal Agency for Support and Evaluation of Graduate Education (CAPES-Brazil) to PVLS.

## Author Contributions

S.T. conceived the project. S.T., and D.M.D., designed the research. S.T., P.G., D.M.D., and A.R.F. supervised the project. S.T., N.K., J.N., K.J., K.J., S.A., Y.Z., P.V.L.S., L-Y.H., and D.M.D. performed the research and analyzed data. A.R.F. and M.H. provided experimental equipment and tools. S.T. wrote the article, with additions and revisions from P.G., D.M.D., A.R.F. and M.H. All authors have read and approved the final version of the manuscript.

## Supplemental data

### Tables

**Supplemental Table S1.** Metabolite contents of the TCA-cycle and BCAAs in the mutant set grown under HC conditions. Plants were grown under environmental controlled conditions in HC (3000 ppm CO_2_) to growth stage 5.1 (Boyes et al., 2001) and leaf-material was harvested at the end of the day (EoD, 9 h illumination). Values are means ± SD (n = 6) and bold letters indicate values statistically significant from the wild type based on Students *t*-test (\**p* < 0.05, \*\**p* < 0.01).

| Metabolite | Col.0 | *trxo1-1* | *trxh2-2* | *gldt1-1* |
| --- | --- | --- | --- | --- |
| **TCA-cycle** |  |  |  |  |
| Pyruvate | 11.71 ± 03.04 | 10.64 ± 03.11 | 10.59 ± 02.69 | 11.34 ± 01.48 |
| Citrate | 3509.25 ± 775.89 | **1638.92 ± 354.92\*\*** | 2852.11 ± 844.49 | 3801.61 ± 613.98 |
| Aconitate | 63.48 ± 10.80 | **37.11 ± 06.39\*\*** | **34.62 ± 06.26\*\*** | 60.76 ± 07.07 |
| 2-Oxoglutarate | 15.30 ± 03.46 | **08.82 ± 03.32\*** | 14.53 ± 03.40 | **09.89 ± 01.90\*\*** |
| Succinate | 07.67 ± 02.26 | **10.95 ± 02.43\*** | 9.96 ± 04.12 | 06.63 ± 01.62 |
| Fumarate | 45.00 ± 09.53 | **57.60 ± 10.46\*** | 63.27 ± 13.05 | 42.12 ± 11.81 |
| Malate | 15862.56 ± 2491.80 | 14883.41 ± 1789.77 | 14570.21 ± 1713.66 | 13186.09 ± 2425.34 |
| GABA | 2.68 ± 01.25 | 02.46 ± 01.30 | 04.95 ± 01.70 | 2.59 ± 00.75 |
| **BCAAs** |  |  |  |  |
| Valine | 06.38 ± 00.92 | 06.09 ± 00.97 | 06.56 ± 01.13 | **07.71 ± 01.12\*** |
| Leucine | 01.17 ± 00.23 | 01.23 ± 00.29 | 01.66 ± 00.27 | 01.18 ± 00.48 |
| Isoleucine | 02.04 ± 00.29 | 01.96 ± 00.27 | 02.39 ± 00.41 | **02.74 ± 00.51\*** |
| Metabolite | *trxo1/h2* | *trxo1xgldt1* | *trxh2xgldt1* | *triple* |
| **TCA-cycle** |  |  |  |  |
| Pyruvate | 12.15 ± 01.86 | **16.25 ± 03.87\*** | 14.36 ± 03.53 | **15.06 ± 01.16\*** |
| Citrate | 3876.59 ± 731.59 | 3827.85 ± 1351.45 | **2170.66 ± 634.04\*** | 2836.99 ± 781.77 |
| Aconitate | 56.25 ± 11.31 | 69.07 ± 24.67 | 53.34 ± 14.20 | 55.24 ± 13.18 |
| 2-Oxoglutarate | **10.44 ± 01.78\*** | 17.62 ± 04.99 | 14.62 ± 02.16 | **09.45 ± 01.51\*\*** |
| Succinate | 07.01 ± 02.00 | **13.91 ± 03.63\*** | 11.97 ± 06.23 | 10.60 ± 02.24 |
| Fumarate | 52.52 ± 10.66 | **73.87 ± 19.57\*** | 65.19 ± 14.36 | **57.01 ± 06.65\*** |
| Malate | 14543.40 ± 1755.47 | 17057.54 ± 00.00 | 15144.85 ± 1697.11 | 14446.00 ± 1322.41 |
| GABA | 1.88 ± 01.00 | 01.56 ± 00.71 | 01.53 ± 00.80 | 01.40 ± 00.27 |
| **BCAAs** |  |  |  |  |
| Valine | 06.46 ± 00.73 | 06.67 ± 00.94 | 06.10 ± 01.20 | 06.16 ± 00.82 |
| Leucine | **01.50 ± 00.27\*** | 01.43 ± 00.25 | 01.28 ± 00.24 | **01.58 ± 00.37\*** |
| Isoleucine | 02.12 ± 00.39 | 02.49 ± 00.82 | 02.02 ± 00.47 | 02.16 ± 00.38 |

**Supplemental Table S2.** Metabolite contents of the TCA-cycle and BCAAs in the mutant set grown under HC and shifted to LC conditions. Plants were grown under environmental controlled conditions in HC (3000 ppm CO_2_) to growth stage 5.1 (Boyes et al., 2001) and leaf-material was harvested at the end of the day (EoD, 9 h illumination) following a shift to normal air (LC – 390 ppm CO_2_). Values are means ± SD (n = 6) and bold letters indicate values statistically significant from the wild type based on Students *t*-test (\**p* < 0.05, \*\**p* < 0.01).

| Metabolite | Col.0 | *trxo1-1* | *trxh2-2* | *gldt1-1* |
| --- | --- | --- | --- | --- |
| **TCA-cycle** |  |  |  |  |
| Pyruvate | 14.95 ± 01.64 | 13.42 ± 02.52 | **11.02 ± 01.90\*\*** | **11.03 ± 02.11\*\*** |
| Citrate | 2391.02 ± 319.46 | **3006.46 ± 221.97\*\*** | **1314.60 ± 628.30\*\*** | **1761.96 ± 263.84\*\*** |
| Aconitate | 54.64 ± 11.26 | 65.01 ± 08.70 | 40.77 ± 13.92 | 46.69 ± 04.70 |
| 2-Oxoglutarate | 4.40 ± 01.83 | 3.27 ± 01.44 | 2.86 ± 00.63 | **76.58 ± 29.48\*\*** |
| Succinate | 21.54 ± 05.83 | 19.01 ± 07.28 | 19.57 ± 06.72 | **66.08 ± 12.50\*\*** |
| Fumarate | 69.52 ± 13.95 | 62.21 ± 12.90 | 70.98 ± 13.66 | 69.12 ± 14.18 |
| Malate | 13910.32 ± 1227.38 | 12727.51 ± 1131.78 | 14518.79 ± 2078.32 | 13264.26 ± 2239.30 |
| GABA | 2.51 ± 00.86 | 2.56 ± 00.65 | 2.42 ± 00.63 | 2.71 ± 00.78 |
| **BCAAs** |  |  |  |  |
| Valine | 4.75 ± 00.74 | 5.21 ± 00.93 | 4.87 ± 01.18 | **8.47 ± 02.19\*\*** |
| Leucine | 1.02 ± 00.31 | 1.46 ± 00.78 | 1.55 ± 00.72 | 3.89 ± 01.78 |
| Isoleucine | 1.88 ± 00.48 | 2.20 ± 00.45 | 1.87 ± 00.63 | **5.81 ± 01.48\*\*** |
| **Metabolite** | ***trxo1h2*** | ***trxo1xgldt1*** | ***trxh2xgldt1*** | ***triple*** |
| **TCA-cycle** |  |  |  |  |
| Pyruvate | **19.22 ± 02.34\*\*** | **28.03 ± 04.72\*\*** | **22.19 ± 02.99\*\*** | **19.49 ± 02.08\*\*** |
| Citrate | 2652.83 ± 495.65 | **4969.58 ± 1202.37\*\*** | 2596.87 ± 335.76 | 2491.56 ± 574.39 |
| Aconitate | 51.10 ± 11.79 | **73.53 ± 08.55\*** | **37.06 ± 08.95\*** | 46.61 ± 11.84 |
| 2-Oxoglutarate | **1.81 ± 00.34\*** | **121.59 ± 37.61\*\*** | **110.49 ± 39.12\*\*** | **124.43 ± 26.55\*\*** |
| Succinate | **12.88 ± 04.13\*** | **71.70 ± 16.44\*\*** | **97.19 ± 23.17\*\*** | **101.00 ± 31.86\*\*** |
| Fumarate | 73.10 ± 16.32 | **104.09 ± 26.34\*** | **102.23 ± 22.81\*\*** | **87.57 ± 08.86\*** |
| Malate | 13998.61 ± 1519.49 | **18294.34 ± 3808.52\*\*** | **17847.73 ± 3534.63\*\*** | 14037.86 ± 1020.88 |
| GABA | **0.91 ± 00.12\*\*** | **1.52 ± 00.29\*** | **0.76 ± 00.24\*\*** | **0.96 ± 00.25\*\*** |
| **BCAAs** |  |  |  |  |
| Valine | **3.21 ± 00.15\*\*** | 6.92 ± 03.74 | **8.65 ± 01.22\*\*** | **8.63 ± 01.82\*\*** |
| Leucine | **0.69 ± 00.16\*\*** | **6.79 ± 02.46\*\*** | **6.81 ± 01.51\*\*** | **5.38 ± 01.75\*\*** |
| Isoleucine | **1.16 ± 00.11\*\*** | **7.15 ± 02.27\*\*** | **8.89 ± 02.70\*\*** | **7.95 ± 02.84\*\*** |

**Supplemental Table S3.** Metabolite contents of the TCA-cycle and BCAAs in the mutant set grown under HC conditions. Plants were grown under environmental controlled conditions in LC (390 ppm CO_2_) to growth stage 5.1 (Boyes et al., 2001) and leaf-material was harvested at the end of the day (EoD, 9 h illumination). Values are means ± SD (n = 6) and bold letters indicate values statistically significant from the wild type based on Students *t*-test (\**p* < 0.05, \*\**p* < 0.01).

| Metabolite | Col.0 | *trxo1-1* | *trhx2-2* | *gldt1-1* |
| --- | --- | --- | --- | --- |
| **TCA-cycle** |  |  |  |  |
| Pyruvate | 16.69 ± 02.64 | 16.95 ± 02.16 | **26.94 ± 01.53\*\*** | **33.05 ± 05.59\*\*** |
| Citrate | 1949.32 ± 680.22 | 1755.47 ± 406.63 | 1387.44 ± 445.54 | **4144.19 ± 415.58\*\*** |
| Aconitate | 28.35 ± 07.11 | 28.71 ± 05.15 | **12.05 ± 02.69\*\*** | **40.10 ± 02.58\*** |
| 2-Oxoglutarate | 4.68 ± 01.16 | 5.08 ± 00.98 | **8.00 ± 01.69\*\*** | **212.18 ± 49.85\*\*** |
| Succinate | 6.38 ± 00.70 | **8.56 ± 02.37\*** | **18.18 ± 03.85\*\*** | **77.59 ± 11.76\*\*** |
| Fumarate | 80.60 ± 07.74 | **67.53 ± 11.00\*** | **111.58 ± 26.24\*** | 95.16 ± 20.72 |
| Malate | 8836.46 ± 1094.54 | **10719.37 ± 1144.81\*\*** | **17106.38 ± 2109.96\*\*** | **22456.49 ± 2546.32\*\*** |
| GABA | 0.65 ± 00.12 | 0.68 ± 00.22 | **1.49 ± 00.35\*\*** | **1.19 ± 00.38\*** |
| **BCAAs** |  |  |  |  |
| Valine | 2.17 ± 00.62 | **2.76 ± 00.34\*** | 1.75 ± 00.23 | 2.19 ± 00.18 |
| Leucine | 0.44 ± 00.15 | **0.59 ± 00.09\*** | 0.37 ± 00.06 | **1.33 ± 00.19\*\*** |
| Isoleucine | 0.80 ± 00.25 | **1.04 ± 00.15\*** | 0.79 ± 00.17 | **1.72 ± 00.25\*\*** |
| Metabolite | *trxo1h2* | *trxo1xgldt1* | *trhx2xgldt1* | *triple* |
| **TCA-cycle** |  |  |  |  |
| Pyruvate | **11.54 ± 03.00\*\*** | **28.49 ± 03.32\*\*** | **23.20 ± 03.93\*\*** | **25.26 ± 01.89\*\*** |
| Citrate | 2984.11 ± 905.10 | **4256.00 ± 984.41\*\*** | **4411.76 ± 896.08\*\*** | **5068.81 ± 209.28\*\*** |
| Aconitate | 24.48 ± 04.42 | **47.08 ± 09.04\*\*** | **48.88 ± 07.68\*\*** | **50.72 ± 03.84\*\*** |
| 2-Oxoglutarate | 5.54 ± 01.10 | **385.62 ± 86.18\*\*** | **418.48 ± 81.99\*\*** | **507.66 ± 46.35\*\*** |
| Succinate | **27.02 ± 03.43\*\*** | **45.93 ± 03.80\*\*** | **43.76 ± 06.58\*\*** | **41.91 ± 05.37\*\*** |
| Fumarate | **102.63 ± 18.90\*** | **52.14 ± 13.37\*\*** | **38.29 ± 15.69\*\*** | **22.97 ± 05.32\*\*** |
| Malate | **17271.74 ± 1417.70\*\*** | **22801.14 ± 3476.95\*\*** | **25010.08 ± 4520.13\*\*** | **26233.46 ± 2085.17\*\*** |
| GABA | 1.03 ± 00.35 | **1.04 ± 00.16\*\*** | **1.15 ± 00.20\*\*** | **1.32 ± 00.21\*\*** |
| **BCAAs** |  |  |  |  |
| Valine | 2.09 ± 00.25 | **3.56 ± 00.67\*\*** | **4.26 ± 00.56\*\*** | **4.78 ± 00.86\*\*** |
| Leucine | 0.39 ± 00.11 | **2.17 ± 00.59\*\*** | **2.61 ± 00.53\*\*** | **2.86 ± 00.64\*\*** |
| Isoleucine | 0.67 ± 00.10 | **3.37 ± 00.90\*\*** | **4.09 ± 00.84\*\*** | **4.32 ± 00.99\*\*** |

**Supplemental Table S4.** Pyridine nucleotide contents in the mutant set grown under HC conditions. Plants were grown under environmental controlled conditions in HC (3000 ppm CO_2_) to growth stage 5.1 (Boyes et al., 2001) and leaf-material was harvested at the end of the day (EoD, 9 h illumination). Values are means ± SD (n = 6) and bold letters indicate values statistically significant from the wild type based on Students *t*-test (\**p* < 0.05, \*\**p* < 0.01).

| Genotype | NADH | NAD<sup>+</sup> | NADH/NAD<sup>+</sup> |
| --- | --- | --- | --- |
| Col.0 | 2.49 ± 0.64 | 20.96 ± 2.19 | 0.119 ± 0.008 |
| *trxo1* | 2.90 ± 0.39 | 20.34 ± 2.08 | 0.144 ± 0.027 |
| *trxh2* | 2.75 ± 0.55 | 21.39 ± 2.34 | 0.139 ± 0.031 |
| *gldt1* | 2.78 ± 0.49 | 20.45 ± 1.24 | 0.136 ± 0.025 |
| *trxo1xgldt1* | 2.38 ± 0.46 | 20.98 ± 2.81 | 0.118 ± 0.020 |
| *trxh2xgldt1* | 2.44 ± 0.48 | 19.17 ± 3.17 | 0.128 ± 0.020 |
| *triple* | 2.29 ± 0.26 | **18.03 ± 0.75\*** | 0.130 ± 0.017 |
| *trxo1h2* | 2.38 ± 0.46 | 18.08 ± 2.91 | **0.145 ± 0.010\*\*** |
|  | NADPH | NADP<sup>+</sup> | NADPH/NADP<sup>+</sup> |
| Col.0 | 0.74 ± 0.22 | 14.21 ± 1.49 | 0.052 ± 0.013 |
| *trxo1* | 0.62 ± 0.23 | 13.15 ± 1.92 | 0.047 ± 0.016 |
| *trxh2* | 0.65 ± 0.19 | 11.28 ± 1.30 | 0.057 ± 0.013 |
| *gldt1* | 0.75 ± 0.15 | **11.17 ± 1.63\*\*** | 0.067 ± 0.008 |
| *trxo1xgldt1* | 0.66 ± 0.08 | **11.97 ± 1.59\*** | 0.052 ± 0.007 |
| *trxh2xgldt1* | 0.65 ± 0.13 | **10.76 ± 1.22\*\*** | 0.060 ± 0.010 |
| *triple* | 0.59 ± 0.15 | **10.26 ± 1.35\*\*** | 0.057 ± 0.015 |
| *trxo1h2* | 0.71 ± 0.15 | **9.90 ± 0.60\*\*** | 0.072 ± 0.015 |

**Supplemental Table S5.**
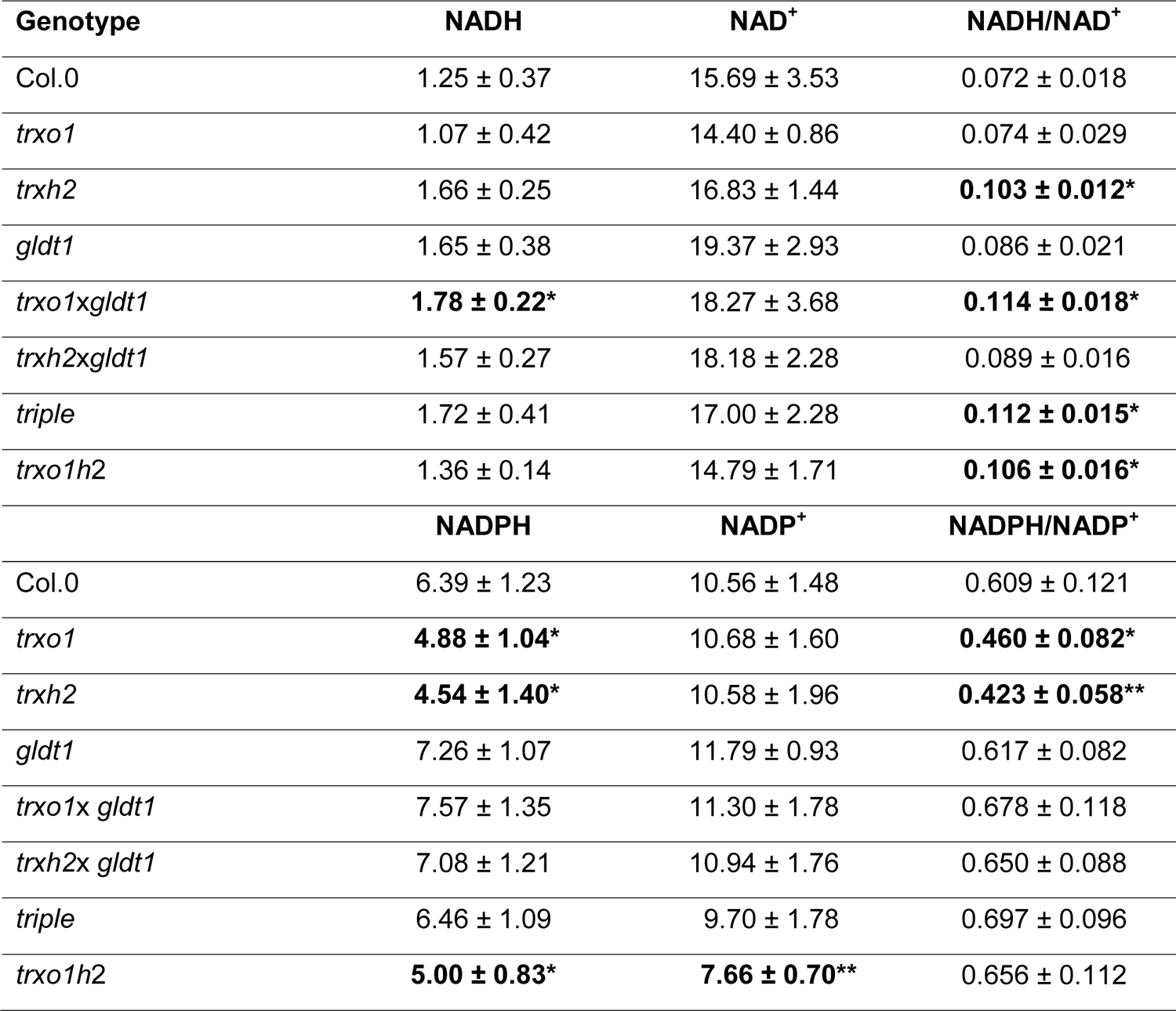
Pyridine nucleotide contents in the mutant set shifted from HC-to-LC conditions. Plants were grown under environmental controlled conditions in HC to growth stage 5.1 (Boyes et al., 2001) and shifted to LC (390 ppm CO_2_). Leaf-material was harvested at the end of the day (EoD, 9 h illumination). Values are means ± SD (n = 6) and bold letters indicate values statistically significant from the wild type based on Students *t*-test (\**p* < 0.05, \*\**p* < 0.01).

**Supplemental Table S6.** Pyridine nucleotide contents in the mutant set under LC conditions. Plants were grown under environmental controlled conditions in LC (390 ppm CO_2_) to growth stage 5.1 (Boyes et al., 2001) and leaf-material was harvested at the end of the day (EoD, 9 h illumination). Values are means ± SD (n = 6) and bold letters indicate values statistically significant from the wild type based on Students *t*-test (\**p* < 0.05, \*\**p* < 0.01).

| Genotype | NADH | NAD<sup>+</sup> | NADH/NAD<sup>+</sup> |
| --- | --- | --- | --- |
| Col.0 | 3.31 ± 0.84 | 24.41 ± 4.27 | 0.136 ± 0.017 |
| *trxo1* | 2.45 ± 0.71 | 22.36 ± 3.90 | 0.108 ± 0.024 |
| *trxh2* | 3.10 ± 0.67 | 27.99 ± 7.29 | 0.116 ± 0.039 |
| *gldt1* | 3.22 ± 0.78 | 28.01 ± 3.48 | 0.108 ± 0.011 |
| *trxo1xgldt1* | 3.23 ± 0.52 | 30.95 ± 4.68 | **0.099 ± 0.001\*\*** |
| *trxh2xgldt1* | 2.97 ± 0.23 | 30.54 ± 2.48 | **0.102 ± 0.010\*\*** |
| *triple* | 2.89 ± 0.66 | 31.23 ± 2.79 | **0.096 ± 0.015\*** |
| *trxo1h2* | 2.66 ± 0.25 | 25.97 ± 1.59 | **0.108 ± 0.017\*** |
|  | NADPH | NADP<sup>+</sup> | NADPH/NADP<sup>+</sup> |
| Col.0 | 0.96 ± 0.18 | 16.40 ± 3.59 | 0.077 ± 0.011 |
| *trxo1* | 0.80 ± 0.24 | **9.73 ± 1.25\*\*** | 0.081 ± 0.018 |
| *trxh2* | 0.93 ± 0.24 | 11.72 ± 1.82 | 0.080 ± 0.023 |
| *gldt1* | 1.15 ± 0.30 | **11.33 ± 1.03\*** | **0.101 ± 0.018\*** |
| *trxo1x gldt1* | **1.62 ± 0.14\*\*** | 15.26 ± 1.51 | **0.109 ± 0.015\*\*** |
| *trxh2x gldt1* | **1.41 ± 0.07\*\*** | **11.83 ± 1.13\*** | **0.129 ± 0.011\*\*** |
| *triple* | **1.39 ± 0.21\*** | 15.05 ± 3.12 | **0.104 ± 0.016\*** |
| *trxo1h2* | 0.94 ± 0.10 | **9.51 ± 1.20\*\*** | **0.100 ± 0.017\*** |

### Figures

**Supplemental Figure S1.**
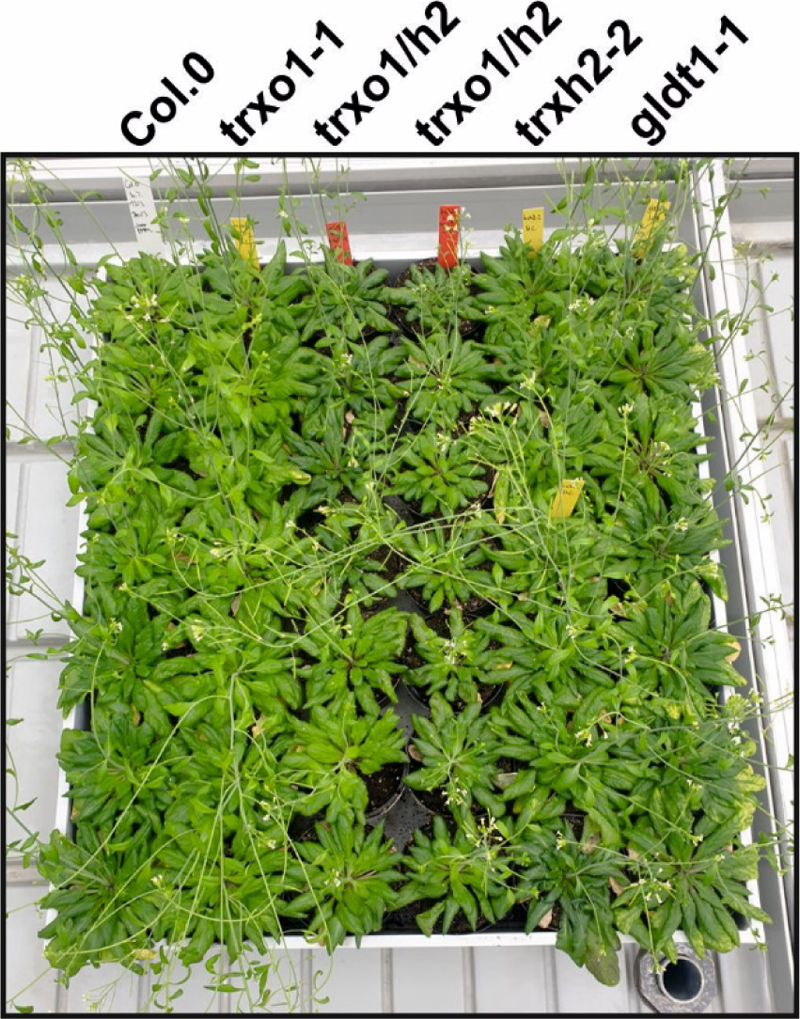
Visual phenotype of *trxo1h2* shifted from HC-to-LC conditions. Plants were grown under standard growth conditions (10/14 h day-/night-cycle) in high CO_2_ (HC, 3000 ppm CO_2_). After reaching growth stage 5.1 (Boyes et al., 2001) plants were transferred to normal air (LC, 390 ppm CO_2_) and growth continued under long day conditions (16/8 h day-/night-cycle) to stimulate flowering and seed production. The picture shown was taken following 2 weeks after the transfer and indicates a severe growth reduction of the *trxo1h2* double mutant under these conditions.

**Supplemental Figure S2.**
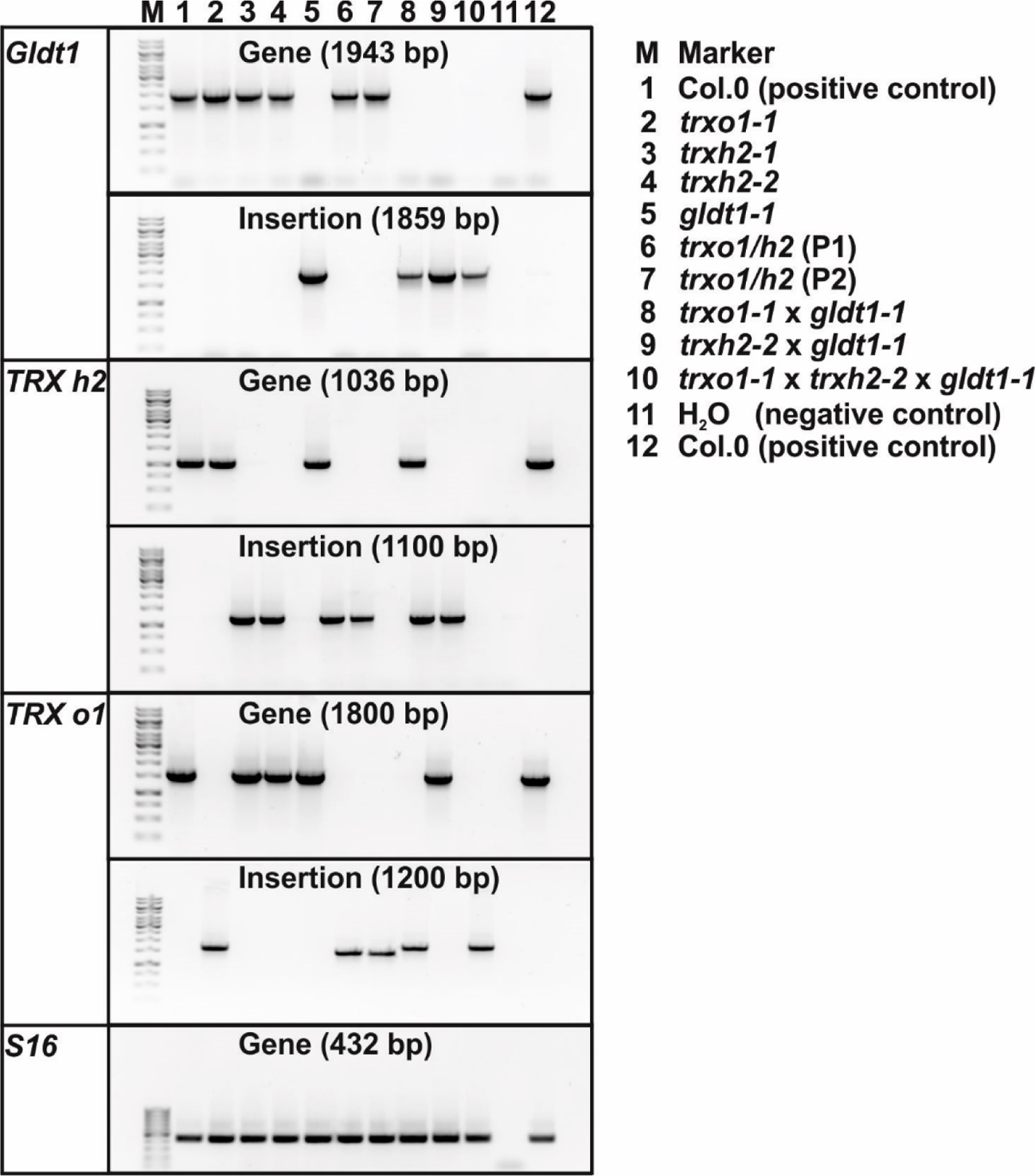
Genotype of multiple *TRX*/*GLDT* mutants. PCR verification of diagnostic fragments for the *GLDT1*, *TRX h2* and *TRX o1* genes and insertions within the respective genes. The *S16* gene was amplified as positive control to verify DNA-integrity.

## Notes

### Competing Interest Statement

The authors have declared no competing interest.

